# Pre-catalytic activation pathway of the Smc5/6 complex

**DOI:** 10.64898/2026.10.09.757879

**Authors:** Jan Siu Kei Wong, Mun Siong Loke, Biswajit Pradhan, Takaharu Kanno, Camilla Björkegren, Eugene Kim

## Abstract

The Smc5/6 complex (Smc5/6) preserves genome stability through ATP-dependent DNA organization, but the structural details that prime the complex for productive DNA engagement remain unknown. Here, we combine cryo-electron microscopy and conformational heterogeneity analysis to define the pre-catalytic activation pathway of Smc5/6. We show that in the absence of ATP and DNA, Smc5/6 exists as a heterogeneous conformational ensemble composed predominantly of two conformations termed Bridge and Cross. In the presence of DNA alone, Smc5/6 adopts a Juxtaposed conformation that brings the ATPase heads into proximity, aligning them for nucleotide engagement. With both ATP and DNA associated, Smc5/6 establishes an Engaged architecture, within which DNA stabilizes its head-associated binding cavity through local remodeling of the complex. Conformational heterogeneity analyses suggest a continuous structural transition between the Cross and Juxtaposed states, while anisotropic network modeling identifies the Bridge–Cross–Juxtaposed–Engaged sequence as the mechanically favored pathway connecting the four experimentally observed conformations. Together, these findings support a pre-catalytic conformational pathway for Smc5/6 activation and provide a structural framework for understanding how conformational heterogeneity gives rise to an activation-competent motor state.

## Introduction

Structural maintenance of chromosomes (SMC) complexes are ATP-powered molecular motors that organize and maintain genomes through dynamic interactions with DNA. In eukaryotes, the three SMC complexes (cohesin, condensin, and Smc5/6) regulate chromosome organization, segregation, DNA repair, and gene expression by coupling ATP binding and hydrolysis to large-scale conformational rearrangements that enable DNA loading, translocation, and DNA loop extrusion^1–6^. Although recent structural and single-molecule studies have substantially advanced our understanding of SMC motor function, how these molecular motors transition from resting states to initiate ATP-dependent DNA transactions remains a fundamental unresolved question.

The Smc5/6 complex plays essential roles in preserving genome stability during DNA replication, homologous recombination, and the cellular response to replication stress^1,7,8^. The hexameric core complex consists of the Smc5 and Smc6 ATPases bridged by the kleisin Nse4, the KITE proteins Nse1 and Nse3, and the SUMO ligase Nse2 (Fig. 1a)^4,9^. Recent single-molecule studies demonstrated that hexameric Smc5/6 functions as an ATP-dependent DNA translocase capable of DNA loop extrusion upon dimerization^4^. Previous biochemical and structural studies have identified multiple DNA-binding regions within Smc5/6, including the ATPase-head and the hinge regions^10–12^. How these DNA interactions and motor activities are coordinated with the ATP- and DNA-dependent conformational changes of Smc5/6, however, remains poorly understood.

**Figure 1:**
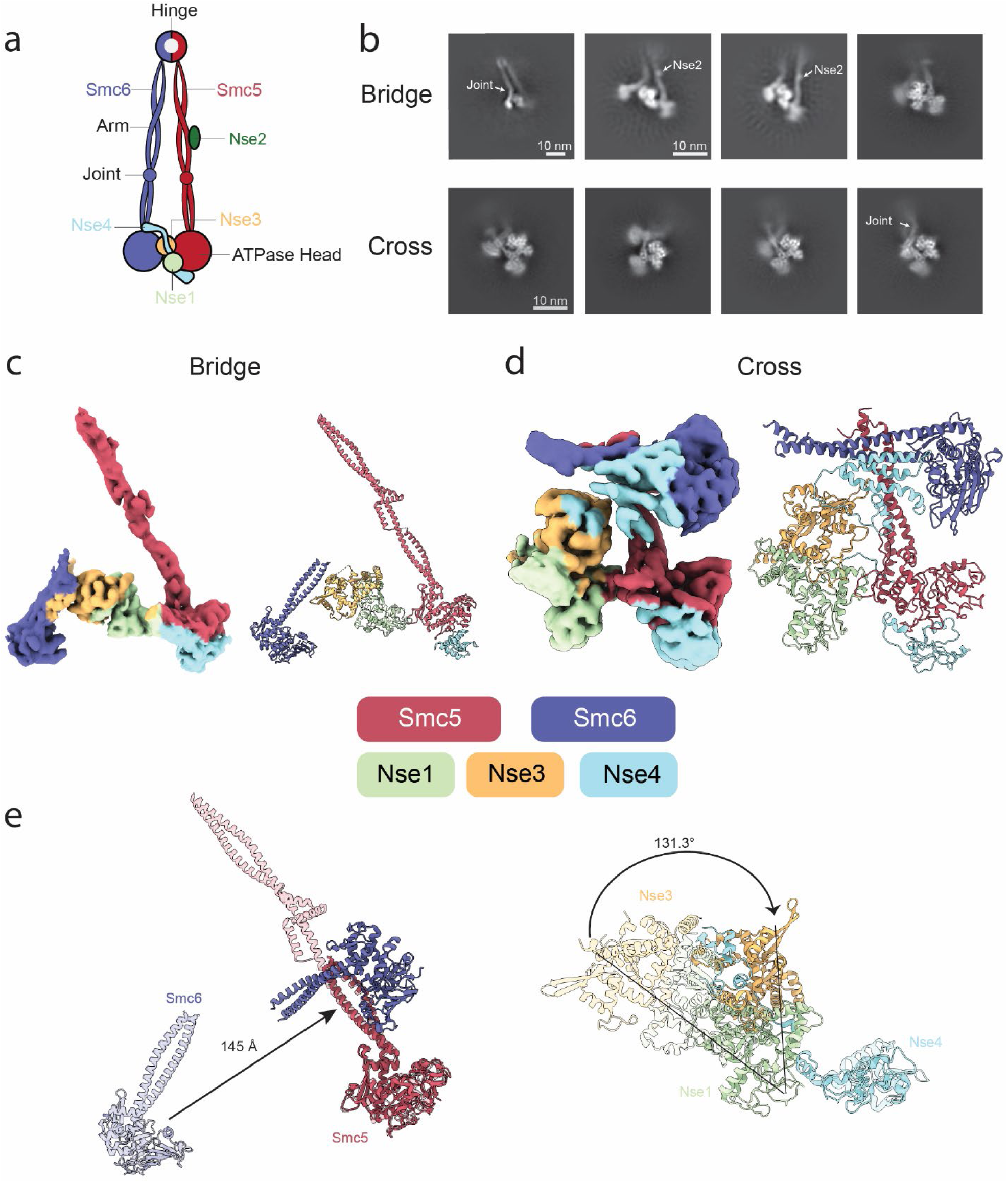
Two distinct apo conformations of the Smc5/6 complex. a, Schematic representation of the *S. cerevisiae* Smc5/6 hexameric core complex. b, Representative 2D class averages of the Bridge and Cross conformations. White arrows indicate bending near the Smc6 joint and Nse2. c,d, Cryo-EM density maps (left) and pseudo-atomic models (right) of the apo Smc5/6 complex in the Bridge (c) and Cross (d) conformations. e, Structural superposition of the Bridge and Cross conformations aligned on the Smc5 coiled coil, highlighting conformational differences in Smc5 and Smc6 (left) and the Nse1–Nse3–Nse4 subcomplex (right).

Recent cryo-electron microscopy (cryo-EM) studies have provided important structural insights into distinct functional states of the Smc5/6 complex. An apo Smc5/6 structure captured in a chemically crosslinked sample adopted a Juxtaposed conformation in which the ATPase heads are brought into close proximity, whereas ATP- and DNA-bound structures captured the ATP-engaged state, in which DNA is accommodated within a head-associated DNA-binding cavity^9,12,13^. Similar Juxtaposed and ATP-engaged architectures have been observed across diverse SMC complexes, suggesting that these configurations represent a conserved mechanochemical state of SMC motors^12–19^. In contrast, atomic force microscopy studies suggested that apo Smc5/6 samples a substantially broader conformational ensemble^20,21^. Whether this apparent structural heterogeneity reflects functionally relevant activation intermediates or simply conformational flexibility has remained unresolved.

Here, we combine cryo-EM and conformational heterogeneity analysis to define the pre-catalytic activation pathway of the Smc5/6 complex. We show that apo Smc5/6 adopts a heterogeneous conformational ensemble that is progressively remodeled toward the ATP-engaged state through distinct structural intermediates. Together, these findings define the structural progression underlying Smc5/6 activation and provide a framework for understanding how conformational heterogeneity gives rise to an activation-competent motor.

### Bridge and Cross are two distinct apo conformations of Smc5/6

To determine the structure of apo Smc5/6, we analyzed the *Saccharomyces cerevisiae* Smc5/6 hexameric core holo-complex by single-particle cryo-EM (Supplementary Fig.1). Two-dimensional (2D) class averages revealed two major conformational states, termed the Bridge and Cross conformations (Fig 1b). Notably, the previously reported juxtaposed conformation observed under chemically cross-linked apo conditions was not observed in our dataset, indicating apo Smc5/6 samples a broader conformational landscape than previously reported^9,13,22,23^.

The Bridge conformation adopts a highly extended architecture, with the ATPase heads separated by approximately 145 Å and connected by the horizontally oriented Nse1/3 subcomplex (Fig. 1c). The Smc5 coiled coil extends in a largely straight trajectory towards the hinge, whereas the Smc6 coiled coil is only partially resolved in the consensus reconstruction. Representative two-dimensional (2D) class averages, however, captured a pronounced bend in the Smc6 arm near the joint region, with substantial variation in the bending angle among different classes (Fig. 1b). This conformational heterogeneity likely explains why only the region of the Smc6 arm below the joint is resolved in the 3D reconstruction.

The Cross conformation adopts a markedly more compact architecture while maintaining disengaged ATPase heads (Fig. 1d). Relative to the Bridge conformation, the SMC6 ATPase head completely crosses over the SMC5 neck, giving rise to the characteristic crossed architecture and reducing the separation between the ATPase heads from approximately 145 Å to 64 Å (Fig. 1e). Concomitantly, the Nse1/3 subcomplex rotates from its horizontal orientation in the Bridge conformation to a more vertical position, nearly parallel to the Smc5 coiled coil (Fig. 1e). This arrangement closely resembles the orientation of Nse1/3 in the previously reported Juxtaposed structure^9,13^. Representative 2D class averages further showed that the pronounced bend in the Smc6 arm is retained in the Cross conformation (Fig. 1b), consistent with substantial intrinsic flexibility of the Smc6 arm.

### DNA incubation selectively enriches the Juxtaposed conformation

Having identified two major apo conformations of Smc5/6, we next asked how DNA influences this conformational ensemble. Cryo-EM analysis of Smc5/6 following incubation with double-stranded DNA yielded a single resolved conformation corresponding to the Juxtaposed state (Fig. 2a–c, Supplementary Fig. 2). The Juxtaposed state is nearly identical to the previously reported chemically cross-linked apo structure, adopting a compact architecture in which the ATPase heads are brought into close proximity and the Smc5 and Smc6 coiled coils become juxtaposed along much of their length (Fig. 2a–c, left)^9,13^. This demonstrates that the Juxtaposed architecture represents an intrinsic conformational state of Smc5/6. Local refinement of the ATPase module enabled reliable modelling of the ATPase heads and associated non-SMC subunits (Fig. 2a–c, right, Supplementary Fig. 2).

**Figure 2:**
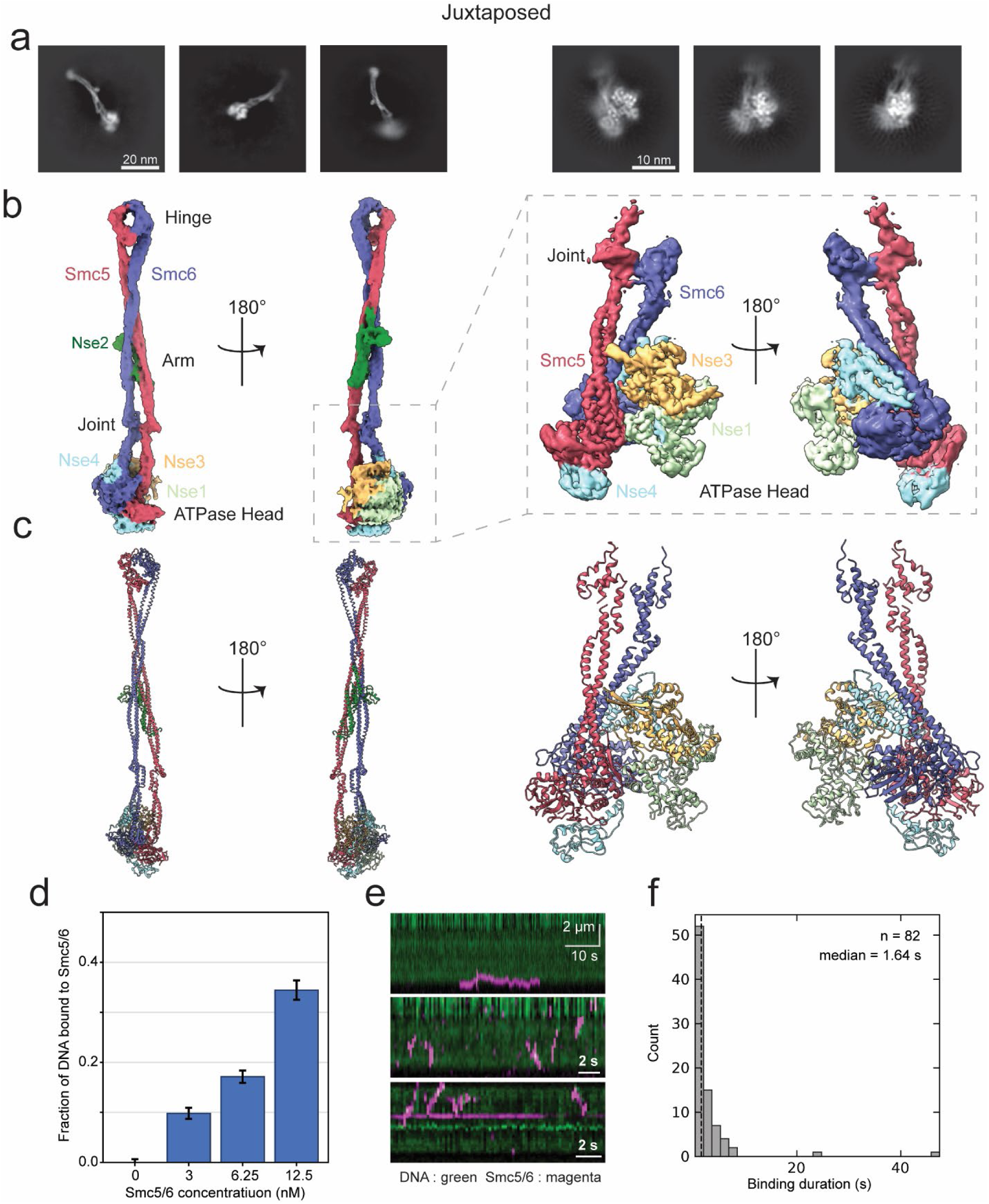
DNA selectively enriches the Juxtaposed conformation. a, Representative 2D class averages of the Juxtaposed conformation. b, Cryo-EM density maps and c, atomic models of the Juxtaposed complex (left) and ATPase-head region (right). d, Fraction of 25 nM Cy5-labeled dsDNA bound as function of Smc5/6 concentration, obtained from Fluorescence correlation spectroscopy (FCS) measurements. e, Representative kymographs of Smc5/6 binding events on DNA in the absence of ATP. f, Distribution of Smc5/6 binding times (n=82) on DNA.

Although DNA was present during sample preparation, no continuous DNA density was observed in the reconstruction. To investigate this apparent discrepancy, we examined ATP-independent DNA binding using fluorescence correlation spectroscopy. Increasing concentrations of Smc5/6 produced a concentration-dependent increase in the DNA-bound fraction, demonstrating that Smc5/6 binds DNA in the absence of ATP (Fig. 2d, Supplementary Fig. 3a-f). Single-molecule imaging similarly captured ATP-independent DNA binding, with kymographs showing frequent association-dissociation of Smc5/6 with DNA (Fig. 2e). Analysis of individual binding events showed that these interactions were short-lived, with residence times of only a few seconds (Fig. 2f). Consistent with these measurements, prolonged incubation of Smc5/6–DNA mixtures at room temperature gradually reduced the DNA-bound fraction, whereas DNA-bound complexes remained stable on ice (Supplementary Fig. 3g).

Together, these observations suggest that transient ATP-independent interactions with DNA shift the Smc5/6 conformational ensemble toward the Juxtaposed state. The short-lived nature of these interactions provides a plausible explanation for the absence of well-resolved DNA density despite the selective enrichment of the Juxtaposed conformation in the presence of DNA.

### Conformational variability links Cross and Juxtaposed states

Comparison of the Cross and Juxtaposed structures showed that the Smc5 ATPase head together with the associated Nse1, Nse3, and Nse4 subunits adopts a highly similar arrangement in both conformations. Alignment using this region as a common reference demonstrated that the principal structural difference lies in the position of the Smc6 ATPase head and the accompanying conformation of the Smc6 coiled coil (Fig. 3a). This observation suggested that the Cross and Juxtaposed states represent distinct positions along a shared conformational trajectory.

**Figure 3:**
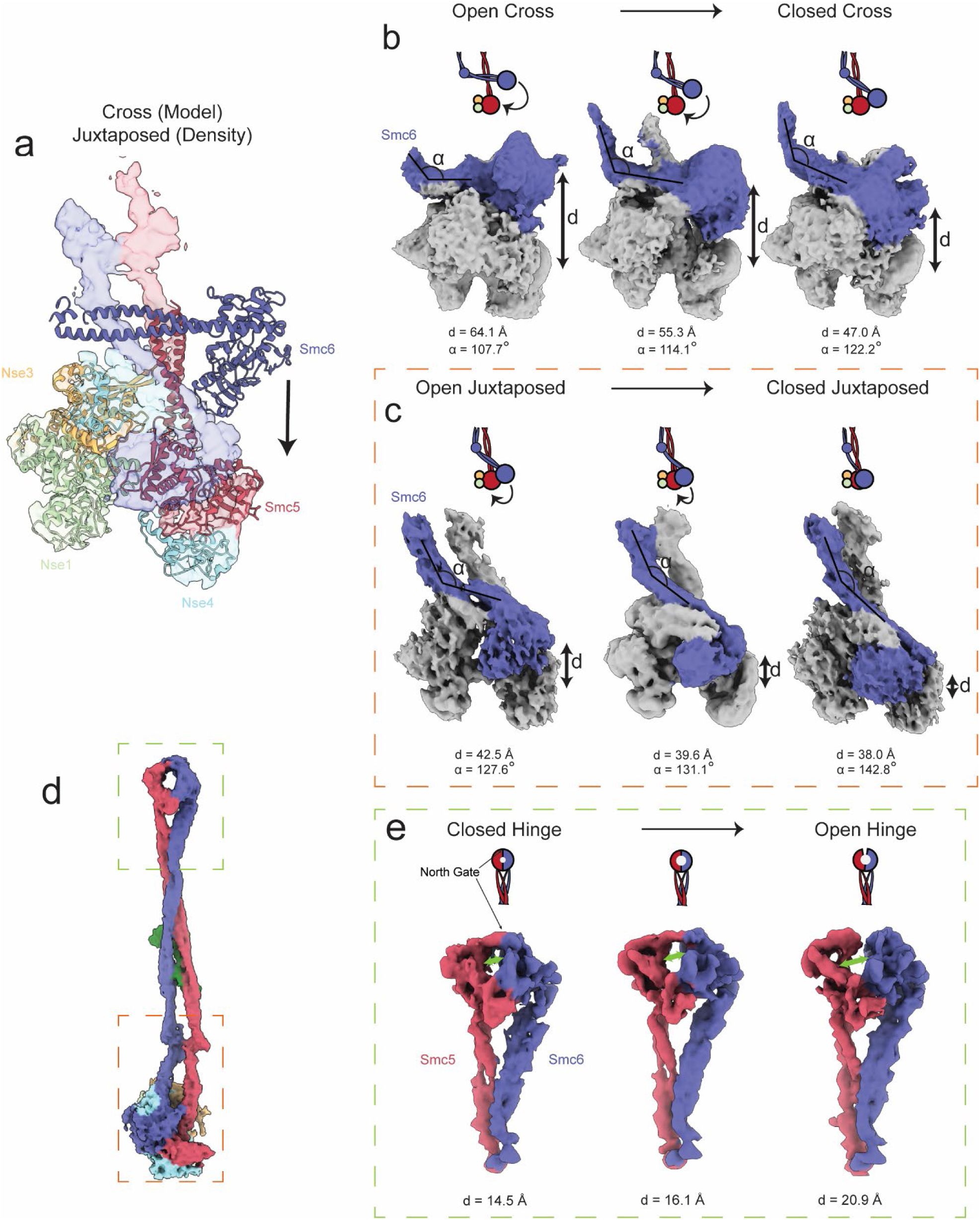
Continuous conformational transition from the Cross to the Juxtaposed state. a, Structural superposition of the Cross and Juxtaposed conformations aligned on the Smc5 coiled coil, showing the conserved arrangement of the Smc5 ATPase head together with the Nse1–Nse3–Nse4 subcomplex. b, cryoDRGN analysis of the Cross ensemble showing progressive movement of the Smc6 ATPase head toward the Smc5 ATPase head, accompanied by increased bending at the Smc6 joint. c, 3D Variability analysis of the Juxtaposed ensemble showing continued approximation of the Smc6 ATPase head toward the Smc5 ATPase head with further bending at the Smc6 joint. d, 3DVA target areas of the Juxtaposed ensemble. e, 3DVA of the hinge in Juxtaposed state showing continuous hinge opening.

To investigate this possibility, we analyzed conformational heterogeneity within the Cross and Juxtaposed particle populations using two complementary approaches cryoDRGN and 3D Variability Analysis (3DVA).^24,25^ Both methods captured similar continuous conformational changes within each ensemble, albeit with differences in structural coverage (Supplementary Fig. 4). In the Cross ensemble, CryoDRGN partially resolved the Smc6 joint region, showing progressive straightening of the Smc6 coiled coil and inward movement of the Smc6 ATPase head toward the Smc5 ATPase head (Fig. 3b, Supplementary Movie 1). Representative conformations along the trajectory showed an increase in the Smc5/6 joint angle from approximately 108° to 122°, accompanied by a reduction in the inter-head distance 64 Å to 47 Å (Fig. 3b). A similar conformational trend was independently observed by 3DVA (Supplementary Fig. 4).

The Juxtaposed ensemble exhibited a continuation of this overall conformational trend. 3DVA revealed further straightening of the Smc6 arm and movement of the Smc6 ATPase head toward the Smc5 ATPase head, with representative conformations showing a reduction in the inter-head distance from 42.5 Å to 38 Å (Fig. 3c, Supplementary Movie 2). CryoDRGN independently captured a similar trend (Supplementary Fig. 4). Notably, toward the most compact conformations of the Cross ensemble, the Smc6 joint angle and inter-head distance approach those observed in the most open conformations of the Juxtaposed ensemble (Fig. 3b, c), further supporting a continuous structural relationship between the two states. Together, these analyses support a continuous structural landscape connecting the Cross and Juxtaposed states, driven primarily by progressive remodeling of the Smc6 arm and repositioning of its ATPase head.

### The Juxtaposed state remains conformationally dynamic at the hinge region

Although the Juxtaposed state adopts a compact overall architecture, local refinement of the hinge and upper coiled-coil region followed by 3DVA revealed substantial conformational dynamics at the hinge (Fig. 3d, Supplementary Fig. 2). The hinge samples a continuous range of closed, intermediate, and open conformations, with its inner diameter increasing from approximately 1.45 nm to 2.1 nm. Notably, the most open conformation exhibited a clear separation of the Smc5 and Smc6 hinge domains at the north interface, which has recently been implicated as a preferential route for ssDNA entry into the hinge lumen^26^. The continuous hinge breathing motion may allow transient capture and release of ssDNA. These observations demonstrate that hinge remains highly dynamic within the compact Juxtaposed conformation, including transient opening of the Smc5–Smc6 interface.

### ATP binding establishes the engaged ATPase architecture, whereas DNA binding induces local remodeling

To determine the structural changes accompanying ATP engagement and DNA binding, we compared Smc5/6 in ATP-bound and ATP/DNA-bound states. The ATP-bound state was captured using a Walker B EQ mutant in the presence of ATP (Fig. 4a, Supplementary Fig. 5), whereas the ATP/DNA-bound state was determined using wild-type Smc5/6 with DNA, ATP, and AMP-PNP (Fig. 4b, Supplementary Fig. 6-8). The ATP-bound structure adopts an engaged ATPase-head architecture closely resembling the previously reported ATP/DNA-bound Smc5/6 structure^12^, demonstrating that ATP binding alone is sufficient to establish the engaged head configuration. Our reconstruction further resolved the conserved β-sheets of both ATPase domains, together with the P-loops and bound nucleotide, providing a more complete structural view of the engaged ATPase module (Supplementary Fig. 7).

**Figure 4:**
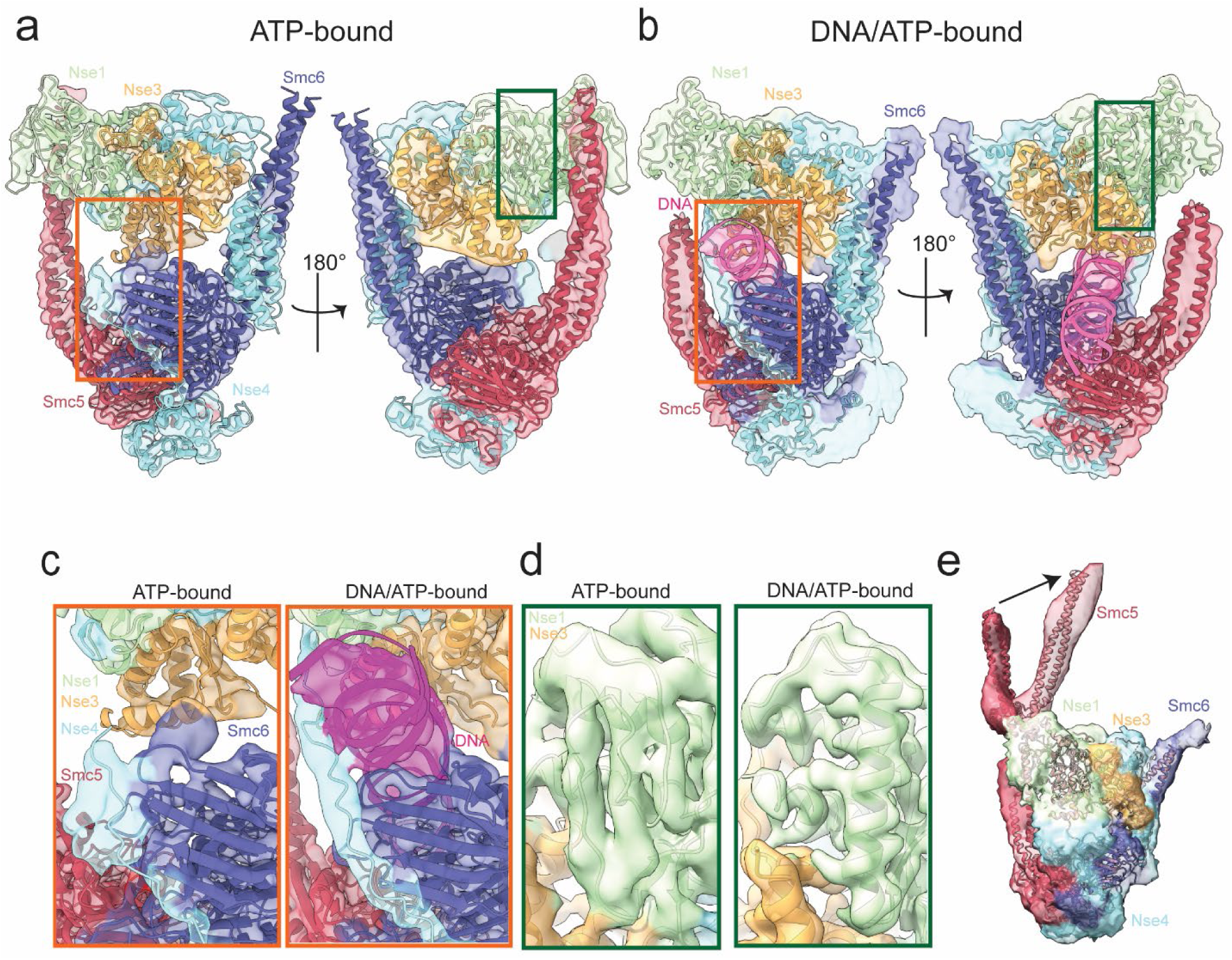
ATP binding establishes the engaged ATPase-head architecture, whereas DNA accommodation induces local remodeling. a, b, Cryo-EM density maps and corresponding atomic models of the ATP-bound Smc5/6 Walker B EQ mutant (a) and the DNA/ATP/AMP-PNP-bound wild-type (WT) complex (b). c, Enlarged view showing interactions between Nse4 and the Smc6 ATPase head in the ATP-bound state Interactions between Nse4 and the Smc6 ATPase head in the ATP-bound state (left) and stabilization of the Nse4 coil around DNA in the DNA/ATP-bound state (right). d, Enlarged view showing DNA-dependent remodeling of Nse1 from a disordered coil in the ATP-bound state (left) to an α-helix in the DNA/ATP-bound state (right). e, CryoDRGN analysis of the ATP-bound Walker B EQ mutant reveals conformational dynamics of the Smc5 coiled-coil arm.

We next examined the conformational heterogeneity within the engaged ATP-bound complex using CryoDRGN. This analysis revealed lateral movement of the Smc5 coiled-coil arm relative to the ATPase-head module, whereas the Smc6 arm remained poorly resolved (Fig. 4e). Thus, substantial conformational flexibility is retained within the complex despite ATP-induced head engagement.

Comparison of the ATP-bound and ATP/DNA-bound structures exhibited only limited structural rearrangements upon DNA binding. Most prominently, Nse4, which is largely flexible in the apo and ATP-bound states, becomes well resolved in the ATP/DNA-bound structure (Fig. 4b). In particular, the Nse4 coil (residues 258–285) forms extensive contacts with the β-sheets at the base of the Smc6 ATPase head, locking the engaged head interface around the bound DNA (Fig. 4c). DNA binding is also accompanied by a subtle repositioning of the Nse1/3 subcomplex and a localized helix-to-coil transition in Nse1 (residues 56–78) adjacent to the DNA-binding cavity (Fig. 4d). The structure further identified new potential DNA-contacting residues, including His56 of Nse3, Val252 of Nse4, Ser137, Ser138, Leu139, and Lys180 of Smc6 (Supplementary Fig. 8).

Taken together, these observations indicate that ATP binding establishes the engaged ATPase architecture of Smc5/6, whereas DNA binding stabilizes and locally remodels the DNA-binding cavity without inducing large-scale structural rearrangements.

### Elastic network modeling supports the sequential activation pathway

Although our structural snapshots define multiple conformational states of Smc5/6, the pathway connecting the apo ensemble to the ATP-engaged architecture remained unclear. Comparison of the four structures revealed a progressive approximation of the ATPase heads during activation (Fig. 5a). The distance between the catalytic domains of Smc5 and Smc6 decreased from 145 Å in the Bridge state to 64 Å in the Cross state, 38 Å in the Juxtaposed state, and 35 Å in the Engaged state. Together with the continuous structural relationship between the Cross and Juxtaposed ensembles (Fig. 3b,c), these observations suggest a potential Bridge → Cross → Juxtaposed → Engaged pathway.

**Figure 5:**
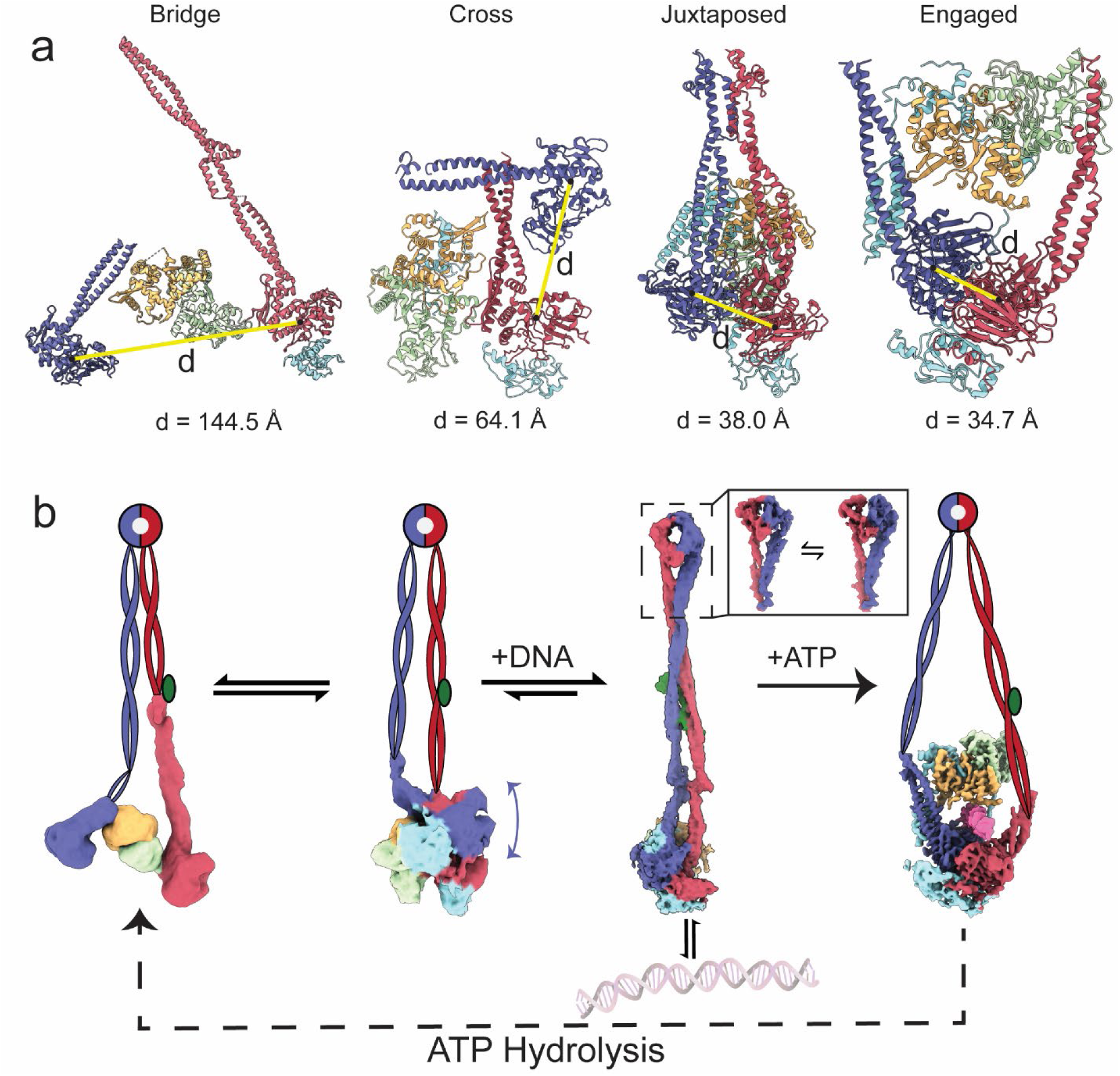
Pre-catalytic activation pathway of the Smc5/6 complex. a, Models of Bridge, Cross, Juxtaposed and Engaged head showing decreasing angles and distances between two ATPase heads. b, Proposed minimum-strain activation pathway connecting the four experimental observed conformations. Under apo condition, Bridge and Cross coexist in a dynamic equilibrium. DNA shifts the conformational ensemble toward the activation-competent Juxtaposed conformation, from which ATP binding establishes the engaged DNA-bound state.

To evaluate whether this sequential pathway is mechanically favoured, we performed anisotropic network model (ANM)-based transition analysis of all 24 possible pathways connecting the four structural states^27^. The Bridge → Cross → Juxtaposed → Engaged pathway consistently minimized deformation energy, ATPase-head displacement, PCA distance, and RMSD (Supplementary Table 1), supporting it as the mechanically favored pathway.

Integrating our structural, biochemical, and computational observations, we propose a sequential activation model for Smc5/6 comprising three distinct transitions (Fig. 5b, Supplementary Movie 3). First, in the absence of ATP, Smc5/6 samples the Bridge and Cross conformations, with the Cross state substantially reducing ATPase-head separation. Second, transient DNA binding in the absence of ATP shifts the conformational equilibrium toward the Juxtaposed state, enriching an activation-competent conformation with closely apposed ATPase heads. Third, ATP binding establishes the Engaged ATPase-head architecture, while DNA binding stabilizes Nse4 within the DNA binding cavity and induces local remodelling of Nse1.

## Discussion

Our findings support a sequential conformational pathway in which Smc5/6 progresses from a dynamic apo ensemble toward ATPase-head engagement through distinct structural intermediates. Rather than undergoing a direct transition between inactive and engaged architectures, Smc5/6 is progressively remodeled through the Bridge, Cross, and Juxtaposed states, with DNA and ATP acting at distinct stages of activation.

### The Cross conformation as a mechanical intermediate

A key finding of this work is the identification of the Cross conformation, a previously unrecognized apo-state intermediate linking the Bridge and Juxtaposed conformations that, to our knowledge, has not been reported in other SMC complexes. Representative 2D classes of the Bridge and Cross conformations indicate that the large-scale rearrangement between these states is accompanied by a pronounced change in Smc6 arm bending around the joint region, from an outward-bent configuration in the Bridge state to an inward-bent configuration in the Cross state. This reversal in bending could facilitate a swing-like motion of the Smc6 arm, allowing its ATPase head to traverse the distance between the Bridge and Cross configurations. Once in the Cross state, conformational heterogeneity analysis suggests that progressive straightening of the inward-bent Smc6 arm further moves its ATPase head toward Smc5, providing a continuous route toward the Juxtaposed state. We therefore propose that the Cross state acts as a mechanical intermediate that couples a joint-mediated swing of the Smc6 arm to its subsequent straightening, thereby progressively bringing the two ATPase heads into proximity. This sequential use of Smc6 joint flexibility may facilitate the substantial architectural transition from the extended Bridge state to the compact, activation-competent Juxtaposed state.

### DNA plays distinct roles before and after ATPase-head engagement

Our findings suggest that DNA plays two distinct roles during Smc5/6 activation. Before ATPase-head engagement, DNA shifts the conformational ensemble toward the Juxtaposed state. Although this conformation was previously captured in chemically crosslinked apo Smc5/6, its selective enrichment by DNA, together with the transient ATP-independent DNA binding observed in our solution and single-molecule measurements, supports its assignment as a functional pre-engaged state. Thus, DNA appears to promote activation through conformational selection of a pre-existing, engagement-competent architecture.

Following ATPase-head engagement, DNA plays a distinct structural role. While ATP establishes the overall Engaged architecture, DNA stabilizes the head-associated DNA-binding cavity through ordering of Nse4 and local remodeling of Nse1. Together, these findings suggest that DNA functions not only as a substrate but also as an allosteric regulator, promoting a pre-engaged conformation before head engagement and stabilizing the DNA-bound architecture thereafter.

### From activation to motor function

Our structures define a pathway leading to ATPase-head engagement but capture only the initial stages of the Smc5/6 mechanochemical cycle. An important outstanding question is whether the conformational states identified here are specific to initial activation or represent recurring intermediates revisited during ATP-driven DNA translocation. In the latter scenario, it is intriguing that the pathway is accompanied by progressive reorganization of the internal architecture of the complex: the Bridge conformation forms a single large chamber, the Cross state partitions this space into two compartments, and the Juxtaposed state closes the upper chamber as the ATPase heads approach, before head engagement establishes two distinct chambers. Such cyclical reorganization could provide a structural basis for coordinating DNA movement through different compartments of the complex. Given the ATP-driven DNA translocation observed in our previous single-molecule studies, determining whether these conformational and topological transitions recur during successive ATPase cycles will be important for understanding how ATP hydrolysis is coupled to DNA translocation and, ultimately, loop extrusion by Smc5/6^4^.

## Data and materials availability

Original imaging data and protein expression constructs are available upon request. The cryo-EM maps and atomic coordinates have been deposited with the Electron Microscopy Data Bank and the PDB with the following accession codes: Apo Bridge (EMD-100183, PDB: 34VB), Apo Cross (EMD-100184, PDB: 34VC), Juxtaposed state overall (EMD-100185, PDB: 34VD), Juxtaposed head (EMD-100186, PDB: 34VE), open Juxtaposed hinge (EMD-100187), intermediate Juxtaposed hinge (EMD-100188), closed Juxtaposed hinge (EMD-100189), +ATP Engaged state (EMD-100190, PDB: 34VF) and +ATP/DNA Engaged state (EMD-100191, PDB: 34VG).

## Code availability

The python script and PDB files contain only Cα atoms for comparing the cumulative deformation energy and structural changes across all possible state orderings are uploaded to GitHub.

## Acknowledgements

We thank Gabriele Maul for protein purification; Sonja Welsch and colleagues for EM support; Melanie McDowell, Noor Agip, Tzu-Jing Yang and Michael Bell for valuable discussion; Scientific Computing Unit and Juan Castillo-Hernandez for IT support.

## Author contributions

C.B. and E.K. initiated the project and acquired funding. J.S.K.W., M.L, T.K., C.B. and E.K. developed the investigation. J.S.K.W., M.L and T.K. purified the Smc5/6 complex. J.S.K.W. and M.L prepared cryoEM grids, collected cryoEM data, processed cryoEM data and built the models. J.S.K.W. and M.L and E.K. analysed and interpreted the models. B.P. collected and analyzed fluorescence correlation spectroscopy data. J.S.K.W. performed elastic network modeling. J.S.K.W., M.L, B.P. and E.K. wrote the manuscript with input from all authors.

## Methods

### Overexpression and purification of Smc5/6

The Smc5/6 complex was overexpressed and purified following a previously established protocol^4^. In brief, full-length Smc5, C-terminal TAP tagged Smc6, Nse1, Nse2, Nse3, and Nse4 were integrated into CB3245. For the ATPase Walker B (EQ) mutant, point mutations were introduced using standard methods at Smc5E1015Q and Smc6E1048Q.

Overexpression strains were grown at 30 °C in 2 L of YEP-lactate medium to optical density (OD600) 0.8–1.0, then protein expression was induced for 4 h by the addition of 2% galactose. After harvesting, cells were disrupted using freezermills (SPEX) with liquid nitrogen cooling, and proteins were extracted by the addition of one cell volume of IPP150 buffer (50 mM Tris-HCl pH 8.0, 150 mM NaCl, 10% glycerol, 0.1% IGEPAL CA-630, 1 mM DTT) containing 10 mM MgCl_2_ and complete EDTA-free protease inhibitor (Roche Applied Science), followed by a treatment with benzonase (Merck) for 1 h at 4 °C. Cleared extracts were mixed with IgG Sepharose 6 FF (Merck) for 2 h at 4 °C and washed with IPP150 buffer. IPP150 buffer was then replaced with GF500 buffer (20 mM HEPES-NaOH pH 7.5, 500 mM NaCl, 10% glycerol, 0.1% IGEPAL CA-630, 1 mM DTT) and the resin treated with tobacco etch virus (TEV) protease at 4 °C overnight with rotation. The fraction eluted by TEV proteinase treatment was diluted four-fold in CBB500 buffer (50 mM Tris-HCl pH 8.0, 500 mM NaCl, 1 mM Mg(CH3COO)_2_, 1 mM imidazole, 2 mM CaCl_2_, 1 mM DTT, 0.1% IGEPAL CA-630) supplemented with 1 M CaCl_2_ (30 µl for 40 ml of mixture) and incubated with calmodulin Sepharose 4B (Merck) for 2 h at 4 °C. After washing with CBB500 buffer, proteins were eluted using CEB500 buffer (50 mM Tris-HCl pH 8.0, 500 mM NaCl, 1 mM Mg(CH3COO)_2_, 1 mM imidazole, 20 mM EGTA, 1 mM DTT, 0.1% IGEPAL CA-630). The eluate was concentrated by around 50-fold using a Vivaspin20 ultrafiltration unit (100 K MWCO, Sartorius) concomitant with an exchange to STO500 buffer (50 mM Tris-HCl pH 8.0, 500 mM NaCl, 2 mM MgCl_2_, 0.5 mM tris(2-carboxyethyl)phosphine (TCEP), 10% glycerol, 0.1% IGEPAL CA-630). Concentration of the complex was determined by Bradford assay using bovine serum albumin (BSA) as standard. The protein was stored at −80 °C at a high concentration.

### Cryo-EM sample preparation and data collection

For Apo, +DNA and ATP-bound states, 3 µL of purified Smc5/6 hexamer WB at a concentration of 100 nM or 0.04 mg/mL with buffer containing 40 mM Tris, pH 7.5, 75 mM - 100mM KGlu, 7.5 mM MgCl_2_ and 1 mM TCEP was applied to freshly glow-discharged Cu grid 2/2 300 mesh (Quantifoil) coated with 1nm continuous carbon layer. The grids were blotted for 6s at 4 °C and 100% humidity and were flash frozen in liquid ethane using an FEI Vitrobot Mark IV (Thermo Fisher).

For ATP/AMP-PNP/DNA-bound state, purified Smc5/6 WT was first diluted to 0.18mg/mL with buffer containing 20 mM HEPES pH 7.5, 2 mM MgCl_2_, 2 % glycerol, 0.5 mM TCEP and 0.06 mM IGEPAL CA-630. The diluted protein was incubated at room temperature for 10 min with 1 mM of ATP, AMP-PNP and a 5-fold excess of 50 bp dsDNA (5’-ATAGTGATATGCGTACGATATAGATCTAGTAGATCGTATGATACGTCGCT-3’, Integrated DNA Technologies, the single-stranded DNA and its complementary DNA were incubated at 95 °C for 5 min and cool down to room temperature at a rate of 1 °C/min). The sample was centrifuged at 14,000 xg for 10 min and the supernatant was transferred to a new tube and kept on ice until the freezing step.

The cryo-electron microscopy grids were washed with 30 min of chloroform bath with magnet stirring and a brief dipping in ethanol before overnight drying. The grids were freshly glow-discharged before sample application (Quantifoil R2/2 Cu 200 mesh grid; Negative gas, 90 s, 15 mA, 15 s holding time, PELCO easiGlow). 3 μl of the DNA-containing protein sample (final concentration of 166 μg/ml) was deposited onto the grid in the chamber of Vitrobot Mark IV (Thermo Fisher Scientific) at 4 °C and 100 % relative humidity, for a period of 10 seconds waiting time, the grids were blotted for 8-9 seconds with 0 blotting force. The grids were immediately plunged into liquid ethane at liquid nitrogen temperature.

Apo-states, +DNA and +ATP/AMP-PNP/DNA-bounds sates micrographs were recorded on a Titan Krios G3i electron microscope (FEI) operated at 300 kV and equipped with a K3 summit direct electron detector (Gatan), mounted behind a GIF Quantum energy filter with a slit width of 30 eV. Images were collected automatically using EPU (Thermo Fisher) in counting mode through twelve separate data collection sessions with a total dose of 60 electrons per Å^2^ during exposure times of 3-3.3 s, dose fractionated into 60 movie frames at defocus ranges of 1.3-2.5 µm. The magnifications used were 105,000×, resulting in physical pixel sizes of 0.837Å.

For the +ATP state, micrographs were recorded on a Titan Krios G2 electron microscope (FEI) operated at 300 kV and equipped with a K3 summit direct electron detector (Gatan), mounted behind a GIF Quantum energy filter with a slit width of 20 eV. A nominal defocus range of –1 to –2.5 µm was used with a total electron dose of 60 electrons per Å^2^ and a pixel size of 0.831 Å per pixel (105,000× magnification) at a tilt angle of 40 °.

### Cryo-EM image processing and 3D reconstruction

All image processing was done in CryoSPARC 4.0-5.0^28^. Movies were aligned using 5 × 5 patches in patch motion correction, and the contrast transfer function (CTF) parameters were estimated with path CTF estimation^29^. Particles were picked manually from a few micrographs, and the resulting particles were binned and extracted with a 168^2^-pixel box size (2 Å per pixel) to generate 2D class averages. Good 2D class averages were used as an initial training set for training the neural-network-based particle pickerTopaz. An initial low-resolution reconstruction was generated using Ab-initio Reconstruction followed by Heterogenous Refinement. Particles of Apo states and ATP-bound state were then re-extracted at 440 pixels with binned pixel to 128. The resulting particle stack was used to retrain the Topaz picker^30^. An iteration comprising particle extraction, 2D classification, Ab-initio reconstruction, Heterogenous Refinement, and Topaz model training was repeated until no further improvement in particle quality was observed. Particles were re-extracted to a binned pixel of 168 then imported to RELION 5 for further refinement with Blush regularization^31^.

To improve alignment of the head domain, volume alignment was performed with a focused on the head region in the +DNA and +ATP/AMP-PNP/DNA state. For +DNA state, particles were re-extracted at 352 pixels with binned pixel to 196. The resulting particle stack was used to retrain the Topaz picker. An iteration comprising particle extraction, 2D classification, Ab-initio reconstruction, Heterogenous Refinement, and Topaz model training was repeated until no further improvement in particle quality was observed. Particles were then imported to RELION for further refinement with Blush regularization, yielding a resolution of 3.8 Å^32^. For +ATP/AMP-PNP/DNA state, particles were re-extracted at 432 pixels. The resulting particle stack was used to retrain the Topaz picker. An iteration comprising particle extraction, 2D classification, Ab-initio reconstruction, Heterogenous Refinement, and Topaz model training was repeated until no further improvement in particle quality was observed. Following the final iteration, a set of 84,394 particles was selected from the best Heterogenous Refinement classes, and was re-extracted at full resolution in an unbinned box of 500 pixels. The particle was subjected to Homogeneous Refinement, yielding an initial resolution of 4.23 Å. With further Non-uniform Refinement and Local Refinement, the resolution was improved to 3.67 Å.

All 3DVA analysis were performed using the refinement mask from the final refinement jobs and a filter resolution of 7 Å for Apo-Cross and 5 Å for Juxtaposed state. For CryoDRGN analysis, particles were binned to 168 pixels with 128 pixels using 3 layers for both input and output layers and 10-dimensional latent space for trainings. All final maps are sharpened using confidence-weighted cryoEM map optimization by variational inference in LocScale2^33^.

### Model building and refinement

Previously published hexameric head structure (PDB:8WJN) and hexameric complex (PDB:8I13) were rigid body fit to EM densities from Apo states and +DNA state in UCSF ChimeraX respectively^9,34^. For different +DNA conformations, the atomic model was refined interactively using ISOLDE v1.9^35^. The refined models were subjected to real-space refinement using the *phenix.real_space_refine* programme implemented in Phenix v1.21-5207^36^. An initial atomic model was generated automatically using ModelAngelo, producing a near-complete starting model for ATP/AMP-PNP/DNA-bound state^37^. The model was imported into UCSF ChimeraX and refined interactively using the ISOLDE v1.11.1. DNA is modelled using AlphaFold 3 according to the known DNA sequence^38^. The model was then further refined in Coot v0.9.8.92^39^. The refined model was subjected to real-space refinement using the *phenix.real_space_refine* programme implemented in Phenix v 1.20.1-4487. The ATP/AMP-PNP/DNA-bound state atomic model was rigid body fit into ATP-bound state density map in UCSF ChimeraX and refined interactively using the ISOLDE v1.9. The refined model was then further refined in Coot v0.9.8.92. The refined model was subjected to real-space refinement using the *phenix.real_space_refine* programme implemented in Phenix v1.21-5207. Molecular graphics and analyses were performed with UCSF ChimeraX.

### DNA-binding assay with Fluorescence correlation spectroscopy (FCS)

DNA binding was assessed by fluorescence correlation spectroscopy using Cy5-labelled 72bp double-stranded DNA at a final concentration of 25 nM. The labelled DNA was mixed with the Smc5/6 complex at final protein concentrations of 0, 3, 6.25, 12.5, 25, 50, 100 or 200 nM in buffer containing 40 mM Tris, pH 7.5, 75mM KGlu, 7.5mM MgCl_2_ and 1mM TCEP. A condition containing 50 nM Smc5/6 complex and 1mM ATP was included as a positive control. Samples were injected into a microfluidic channel whose surfaces were passivated with polyethylene glycol (MW 5000) similar to the single-molecule slides and measured on a Zeiss LSM980 confocal microscope. Measurements were performed at ambient laboratory temperature without active temperature control of the sample stage. FCS and corresponding intensity traces were acquired every 2 s until 300 s after loading of protein-DNA sample.

FCS data were analysed using PyCorrFit v1.1.7. To prevent correlation curves form being skewed by aggregates, curves containing irregular high-intensity bursts were discarded before computing the mean and standard deviation of the remaining FCS. Averaged autocorrelation curves were fitted over lag-time intervals by Levenberg–Marquardt minimisation of the reduced weighted sum of squares, with weights derived from the standard deviation of the averaged curves. The structure parameter SP = z₀/r₀ was fixed to 5.0 as determined from calibration with a free-dye standard of known diffusion coefficient, and the offset was fixed to zero.

Reference samples were fitted with a single-component three-dimensional diffusion model including one blinking (triplet-like) term,

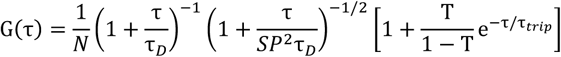

where N is the effective number of particles in the confocal volume, τ_D_ the diffusion time, and T and τ_trip_ the fraction and relaxation time of the non-fluorescent state. Cy5-labelled DNA alone was well described by this single-component model, yielding τ_free_ = 340 µs with τ_trip_ in the range 4–10 µs. In the presence of 50 nM Smc5/6 and 1 mM ATP, a single component was likewise sufficient and yielded τ_bound = 850 µs. Independently, Alexa647-labelled Smc5/6 (via a SNAP-tag on Nse4) measured alone gave a comparable diffusion time of 810 µs, consistent with the 850 µs species corresponding to DNA in complex with Smc5/6.

Autocorrelation curves of the concentration series were then fitted with a two-component three-dimensional diffusion model including one blinking term,

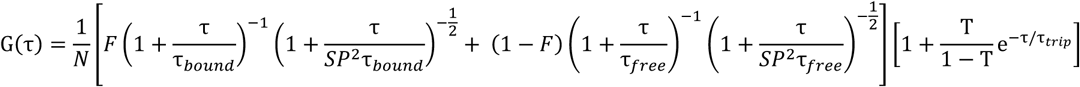

in which the two diffusion times were fixed to the values obtained from the reference measurements (τ_bound_ = 850 µs, τ_free_ = 340 µs) and N, F, T and τ_trip_ were treated as free parameters. The relative molecular brightness of the two species was fixed at α = q_bound_/q_free_ = 1, i.e. the Cy5 fluorophore was assumed to be equally bright in the free and protein-bound DNA, so that F corresponds directly to the fraction of DNA bound to Smc5/6. The bound fraction was determined independently per protein concentration.

### Single molecule imaging of Smc5/6 on λ-DNA

Single-molecule visualization of λ-DNA and labelled Smc5/6 was performed on a cutom-built highly inclined and laminated optical sheet (HILO) microscopy to as described previously^11^. Briefly, 561 nm and 638 nm laser lines were combined into a single-mode fibre and illuminated to the sample through a 100×/1.46 NA oil-immersion TIRF objective at an adjustable illumination angle, which restricted excitation to a thin sheet near the surface and thereby suppressed out-of-focus background and photobleaching. Emission was separated with a multidichroic, residual excitation light removed with a multiband notch filter, and images recorded on an sCMOS camera. Alternating excitation between the two lines was achieved by electronic triggering from an Arduino. The flow cell was held at 30 °C using a resistive heating foil bonded to the slide. Acquisition was controlled by custom Python software; images were recorded at 100 ms exposure per frame for 1000 s.

Flow cells were prepared as described^11^. Glass slides were cleaned in piranha solution, amino-silanised, and passivated with a mixture of methoxy-PEG-NHS (MW 5000) and biotin-PEG-NHS (MW 5000) in borate buffer, with the PEGylation step repeated five times to suppress non-specific protein adsorption. Assembled channels were incubated with 100 nM streptavidin, washed, and 10 pM doubly biotinylated 48.5 kb λ-DNA was flowed in at 3 µl min⁻¹, yielding surface-tethered molecules with relative extensions of 0.1–0.6; unbound DNA was washed out and the channel further blocked with 0.5 mg ml⁻¹ BSA.

DNA-binding experiments were performed in imaging buffer containing 40 mM Tris-HCl pH 7.5, [100 mM NaCl / potassium glutamate], 7.5 mM MgCl₂, 0.5 mg ml⁻¹ BSA, 1 mM TCEP and 200 nM Sytox Orange, supplemented with an oxygen-scavenging and triplet-quenching system (1% D-glucose, 37.5 µM glucose oxidase, 10 nM catalase, 2 mM Trolox). ATP was omitted. Buffer containing 2 nM Alexa 647-labelled Smc5/6 (EQ mutant) was introduced at 30 µl min⁻¹ for 1 min and flow was then stopped. Sytox Orange-stained DNA and labelled Smc5/6 were imaged by alternating excitation at 561 nm (0.1 W cm⁻²) and 638 nm (∼150 W cm⁻²).

Image sequences were analysed with custom Python software (LEADS, https://github.com/biswajitSM/LEADS) ^40^. Regions containing individual λ-DNA molecules were selected manually and cropped; background was removed with a white top-hat filter and a 3-pixel median filter applied before quantification. Kymographs were constructed by summing fluorescence intensity across 11 pixels along lines centred on the DNA axis so that each vertical line corresponds to one frame. Binding events were identified manually and their durations measured from the appearance to the disappearance of the Smc5/6 signal on DNA. Because tethered λ-DNA undergoes transverse thermal fluctuations, DNA-bound complexes could be distinguished from complexes adsorbed to the glass surface by whether the Smc5/6 signal fluctuated together with the underlying DNA; signals that remained stationary independently of the DNA were classified as surface-bound and excluded from analysis (Fig. 2e, last kymograph at the middle) . These surface-adsorbed molecules also served as an in situ measure of the photobleaching lifetime under identical illumination: their fluorescence persisted more than tenfold longer than the measured Smc5/6 binding times on DNA (median 1.64 s), indicating that photobleaching contributes negligibly to the reported binding-time distribution.

### ANM-based elastic deformation analysis of reaction sequence

To compare the relative mechanical resistance among the four experimentally determined conformational states (Bridge, X, J, and E), anisotropic network models (ANMs) consisting exclusively of Cα atoms were generated using ProDy. To establish a common Cartesian reference frame and eliminate rigid-body translation and rotation, states X, J, and E were structurally superposed onto the Bridge state using an identical residue selection. Before superposition, Nse4 and additional residues absent from one or more models were removed to ensure that all structures contained the same set and number of residues. For each directed transition from an initial state i to a target state j, an ANM was constructed using the contact topology of the initial state. Pairs of Cα atoms separated by less than 15.0 Å were connected by identical harmonic springs with a uniform spring constant of *γγ* = 1. The resulting 3N x 3N Hessian matrix (**H_i_**) constructed in ProDy represented the second derivatives of the harmonic ANM potential with respect to with respect to the Cartesian coordinates of the initial state. The Cartesian deformation vector for each directed transition was defined as:

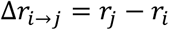

where *r_i_* and *r_jj_* denote the aligned Cartesian coordinates of the common Cα atoms in the initial and target states, respectively. Within the second-order harmonic approximation, the multidimensional form of the Hookean potential was calculate^41^:

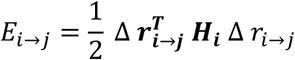

Because the Hessian matrix is constructed independently from the topology of each initial state, the analysis is intrinsically directional (*E_i_*_→*j*_ ≠ *E_j_*_→*i*_ ). For a given candidate sequence (*S*) = (*s*_1_, *s*_2_, *s*_3_, *s*_4_), the cumulative elastic deformation score was defined as:

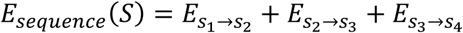

With *γ* = 1, the resulting cumulative score represents the arbitrary harmonic penalty associated with imposing the experimentally observed displacements onto the unrelaxed ANM networks of the initial states. Consequently, these sequence scores serve strictly as relative measures of mechanical compatibility and structural strain. They are not interpreted as thermodynamic free-energy differences, kinetic rates, or physical activation barriers.

The geometric displacement associated with each sequence was independently quantified using three orthogonal metrics: cumulative inter-domain distance fluctuation, cumulative Principal Component Analysis (PCA) distance, and cumulative root-mean-square deviation (RMSD). The inter-domain distance was defined as the scalar distance between the two ATPase heads. PCA was performed on the aligned Cartesian coordinates of the four states to extract the dominant modes of variance, with 2D pairwise distances calculated as Euclidean distances within the PC1-PC2 subspace:

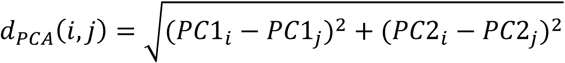

For each candidate state ordering, the cumulative geometric trajectories were calculated by summing the respective absolute variations across the three transitions:

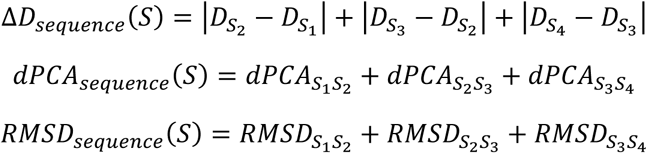

Candidate sequences were ranked primarily by their minimized cumulative ANM deformation score. The cumulative distance fluctuation, PCA distance, and RMSD were evaluated as orthogonal geometric descriptors to validate the physical viability of the spatial trajectories. Sequences exhibiting the simultaneous minimization of cumulative elastic deformation energy and all three geometric distance metrics were determined to be the most mechanically and physically optimal pathways connecting the four observed conformations.

## References

1. Kim, E., Barth, R., and Dekker, C. (2023). Looping the Genome with SMC Complexes. Annu Rev Biochem 92, 15–41. 10.1146/annurev-biochem-032620-110506.

2. Gruber, S. (2017). Shaping chromosomes by DNA capture and release: gating the SMC rings. Curr Opin Cell Biol 46, 87–93. 10.1016/j.ceb.2017.04.002.

3. Ganji, M., Shaltiel, I.A., Bisht, S., Kim, E., Kalichava, A., Haering, C.H., and Dekker, C. (2018). Real-time imaging of DNA loop extrusion by condensin. Science 360, 102–105. 10.1126/science.aar7831.

4. Pradhan, B., Kanno, T., Umeda Igarashi, M., Loke, M.S., Baaske, M.D., Wong, J.S.K., Jeppsson, K., Björkegren, C., and Kim, E. (2023). The Smc5/6 complex is a DNA loop-extruding motor. Nature C*1C*, 843–848. 10.1038/s41586-023-05963-3.

5. Davidson, I.F., Bauer, B., Goetz, D., Tang, W., Wutz, G., and Peters, J.-M. (2019). DNA loop extrusion by human cohesin. Science 3CC, 1338–1345. 10.1126/science.aaz3418.

6. Kim, Y., Shi, Z., Zhang, H., Finkelstein, I.J., and Yu, H. (2019). Human cohesin compacts DNA by loop extrusion. Science 3CC, 1345–1349. 10.1126/science.aaz4475.

7. Peng, X.P., and Zhao, X. (2023). The multi-functional Smc5/6 complex in genome protection and disease. Nat Struct Mol Biol 30, 724–734. 10.1038/s41594-023-01015-6.

8. Rutkauskas, M., and Kim, E. (2025). In vitro dynamics of DNA loop extrusion by structural maintenance of chromosomes complexes. Curr Opin Genet Dev 90, 102284. 10.1016/j.gde.2024.102284.

9. Li, Q., Zhang, J., Haluska, C., Zhang, X., Wang, L., Liu, G., Wang, Z., Jin, D., Cheng, T., Wang, H., et al. (2024). Cryo-EM structures of Smc5/6 in multiple states reveal its assembly and functional mechanisms. Nat Struct Mol Biol 31, 1532–1542. 10.1038/s41594-024-01319-1.

10. Roy, M.-A., Dhanaraman, T., and D’Amours, D. (2015). The Smc5-Smc6 heterodimer associates with DNA through several independent binding domains. Sci Rep 5, 9797. 10.1038/srep09797.

11. Alt, A., Dang, H.Q., Wells, O.S., Polo, L.M., Smith, M.A., McGregor, G.A., Welte, T., Lehmann, A.R., Pearl, L.H., Murray, J.M., et al. (2017). Specialized interfaces of Smc5/6 control hinge stability and DNA association. Nat Commun 8, 14011. 10.1038/ncomms14011.

12. Yu, Y., Li, S., Ser, Z., Kuang, H., Than, T., Guan, D., Zhao, X., and Patel, D.J. (2022). Cryo-EM structure of DNA-bound Smc5/6 reveals DNA clamping enabled by multi-subunit conformational changes. Proc Natl Acad Sci U S A 119, e2202799119. 10.1073/pnas.2202799119.

13. Hallett, S.T., Campbell Harry, I., Schellenberger, P., Zhou, L., Cronin, N.B., Baxter, J., Etheridge, T.J., Murray, J.M., and Oliver, A.W. (2022). Cryo-EM structure of the Smc5/6 holo-complex. Nucleic Acids Res 50, 9505–9520. 10.1093/nar/gkac692.

14. Shi, Z., Gao, H., Bai, X.-C., and Yu, H. (2020). Cryo-EM structure of the human cohesin-NIPBL-DNA complex. Science 368, 1454–1459. 10.1126/science.abb0981.

15. Lee, B.-G., Merkel, F., Allegretti, M., Hassler, M., Cawood, C., Lecomte, L., O’Reilly, F.J., Sinn, L.R., Gutierrez-Escribano, P., Kschonsak, M., et al. (2020). Cryo-EM structures of holo condensin reveal a subunit flip-flop mechanism. Nat Struct Mol Biol 27, 743–751. 10.1038/s41594-020-0457-x.

16. Lee, B.-G., Rhodes, J., and Löwe, J. (2022). Clamping of DNA shuts the condensin neck gate. Proc Natl Acad Sci U S A 119, e2120006119. 10.1073/pnas.2120006119.

17. Bürmann, F., Clifton, B., Koekemoer, S., Wilkinson, O.J., Kimanius, D., Dillingham, M.S., and Löwe, J. (2025). Mechanism of DNA capture by the MukBEF SMC complex and its inhibition by a viral DNA mimic. Cell 188, 2465–2479.e14. 10.1016/j.cell.2025.02.032.

18. Hassler, M., Shaltiel, I.A., Kschonsak, M., Simon, B., Merkel, F., Thärichen, L., Bailey, H.J., Macošek, J., Bravo, S., Metz, J., et al. (2019). Structural Basis of an Asymmetric Condensin ATPase Cycle. Mol Cell 74, 1175–1188.e9. 10.1016/j.molcel.2019.03.037.

19. Shaltiel, I.A., Datta, S., Lecomte, L., Hassler, M., Kschonsak, M., Bravo, S., Stober, C., Ormanns, J., Eustermann, S., and Haering, C.H. (2022). A hold-and-feed mechanism drives directional DNA loop extrusion by condensin. Science 376, 1087–1094. 10.1126/science.abm4012.

20. Umeda, K., Kurokawa, Y., Kodera, N., and Murayama, Y. (2026). Submolecular video-imaging of the Smc5/6 complex topologically bound to DNA. Nat Commun 17, 10398. 10.1038/s41467-026-77122-x.

21. Irvin, E.M., Park, S.T., Miller-Browne, V., Lever, R., Li, S., Waterman, G., Kaur, P., Zhao, X., and Wang, H. (2026). Smc5/6 uses its head-NSE module to preferentially associate with ssDNA gaps and ss-dsDNA junctions. Nucleic Acids Res 54, gkag624. 10.1093/nar/gkag624.

22. Hallett, S.T., Schellenberger, P., Zhou, L., Beuron, F., Morris, E., Murray, J.M., and Oliver, A.W. (2021). Nse5/6 is a negative regulator of the ATPase activity of the Smc5/6 complex. Nucleic Acids Res 49, 4534–4549. 10.1093/nar/gkab234.

23. Serrano, D., Cordero, G., Kawamura, R., Sverzhinsky, A., Sarker, M., Roy, S., Malo, C., Pascal, J.M., Marko, J.F., and D’Amours, D. (2020). The Smc5/6 Core Complex Is a Structure-Specific DNA Binding and Compacting Machine. Mol Cell 80, 1025–1038.e5. 10.1016/j.molcel.2020.11.011.

24. Punjani, A., and Fleet, D.J. (2021). 3D variability analysis: Resolving continuous flexibility and discrete heterogeneity from single particle cryo-EM. J Struct Biol 213, 107702. 10.1016/j.jsb.2021.107702.

25. Zhong, E.D., Bepler, T., Berger, B., and Davis, J.H. (2021). CryoDRGN: reconstruction of heterogeneous cryo-EM structures using neural networks. Nat Methods 18, 176–185. 10.1038/s41592-020-01049-4.

26. Liu, H.W., Roisné-Hamelin, F., Taschner, M., Collier, J., Srinivasan, M., and Gruber, S. (2025). The SMC Hinge is a Selective Gate for Obstacle Bypass. Nat Commun 16, 10457. 10.1038/s41467-025-65408-5.

27. Zhang, S., Krieger, J.M., Zhang, Y., Kaya, C., Kaynak, B., Mikulska-Ruminska, K., Doruker, P., Li, H., and Bahar, I. (2021). ProDy 2.0: increased scale and scope after 10 years of protein dynamics modelling with Python. Bioinformatics 37, 3657–3659. 10.1093/bioinformatics/btab187.

28. Punjani, A., Rubinstein, J.L., Fleet, D.J., and Brubaker, M.A. (2017). cryoSPARC: algorithms for rapid unsupervised cryo-EM structure determination. Nat Methods 14, 290–296. 10.1038/nmeth.4169.

29. Rubinstein, J.L., and Brubaker, M.A. (2015). Alignment of cryo-EM movies of individual particles by optimization of image translations. J Struct Biol 192, 188–195. 10.1016/j.jsb.2015.08.007.

30. Bepler, T., Morin, A., Rapp, M., Brasch, J., Shapiro, L., Noble, A.J., and Berger, B. (2019). Positive-unlabeled convolutional neural networks for particle picking in cryo-electron micrographs. Nat Methods 16, 1153–1160. 10.1038/s41592-019-0575-8.

31. Kimanius, D., Jamali, K., Wilkinson, M.E., Lövestam, S., Velazhahan, V., Nakane, T., and Scheres, S.H.W. (2024). Data-driven regularization lowers the size barrier of cryo-EM structure determination. Nat Methods 21, 1216–1221. 10.1038/s41592-024-02304-8.

32. Kimanius, D., Forsberg, B.O., Scheres, S.H., and Lindahl, E. (2016). Accelerated cryo-EM structure determination with parallelisation using GPUs in RELION-2. Elife 5, e18722. 10.7554/eLife.18722.

33. 33. Bharadwaj, A., de Bruin, R., and Jakobi, A.J. (2026). Confidence-guided cryo-EM map optimisation with LocScale-2.0. Nat Commun. 10.1038/s41467-026-75327-8.

34. Meng, E.C., Goddard, T.D., Pettersen, E.F., Couch, G.S., Pearson, Z.J., Morris, J.H., and Ferrin, T.E. (2023). UCSF ChimeraX: Tools for structure building and analysis. Protein Sci 32, e4792. 10.1002/pro.4792.

35. Croll, T.I. (2018). ISOLDE: a physically realistic environment for model building into low-resolution electron-density maps. Acta Crystallogr D Struct Biol 74, 519–530. 10.1107/S2059798318002425.

36. Afonine, P.V., Poon, B.K., Read, R.J., Sobolev, O.V., Terwilliger, T.C., Urzhumtsev, A., and Adams, P.D. (2018). Real-space refinement in PHENIX for cryo-EM and crystallography. Acta Crystallogr D Struct Biol 74, 531–544. 10.1107/S2059798318006551.

37. Jamali, K., Käll, L., Zhang, R., Brown, A., Kimanius, D., and Scheres, S.H.W. (2024). Automated model building and protein identification in cryo-EM maps. Nature 628, 450–457. 10.1038/s41586-024-07215-4.

38. Abramson, J., Adler, J., Dunger, J., Evans, R., Green, T., Pritzel, A., Ronneberger, O., Willmore, L., Ballard, A.J., Bambrick, J., et al. (2024). Accurate structure prediction of biomolecular interactions with AlphaFold 3. Nature 630, 493–500. 10.1038/s41586-024-07487-w.

39. Emsley, P., Lohkamp, B., Scott, W.G., and Cowtan, K. (2010). Features and development of Coot. Acta Crystallogr D Biol Crystallogr CC, 486–501. 10.1107/S0907444910007493.

40. 40. Pradhan, B., Barth, R., Kim, E., Davidson, I.F., Bauer, B., van Laar, T., Yang, W., Ryu, J.-K., van der Torre, J., Peters, J.-M., et al. (2022). SMC complexes can traverse physical roadblocks bigger than their ring size. Cell Rep 41, 111491. 10.1016/j.celrep.2022.111491.

41. Zheng, W., Brooks, B.R., and Hummer, G. (2007). Protein conformational transitions explored by mixed elastic network models. Proteins CS, 43–57. 10.1002/prot.21465.

