## Supplementary material for "Pre-catalytic activation pathway of the Smc5/6 complex": SI: SI.pdf

**Supplementary Table 1. Elastic network modeling.** Comparison of cumulative deformation energy and structural changes across all possible state orderings. Each row represents one of the 24 possible orderings of the four conformational states (Bridge state: B, Cross state: X, Juxtaposed state: J, and Engaged state: E). For an ordering S1→S2→S3→S4, the cumulative ANM deformation energy, cumulative change in inter-head distance, cumulative PCA distance and cumulative Ca RMSD were calculated. Lower cumulative values represent sequences requiring smaller modelled mechanical deformation or geometric changes.

| Reaction Sequence | Accumulated Distance Fluctuation of ATPase Domains | PCA Distance | Total RMSD | Accumulate Deformation Energy |
| --- | --- | --- | --- | --- |
| B -> X -> J -> E | 109.79 | 115.92 | 144.75 | 21439912.64 |
| B -> X -> E -> J | 113.02 | 138.72 | 153.70 | 24283877.88 |
| B -> E -> J -> X | 139.12 | 134.59 | 156.81 | 24618208.92 |
| X -> B -> E -> J | 193.47 | 146.16 | 161.70 | 25175079 |
| B -> E -> X -> J | 165.22 | 142.85 | 161.60 | 25294191.8 |
| J -> X -> B -> E | 216.34 | 131.63 | 157.55 | 26178592.17 |
| X -> B -> J -> E | 190.24 | 152.33 | 166.12 | 26313092.42 |
| E -> J -> X -> B | 109.79 | 115.92 | 144.75 | 26768366.93 |
| B -> J -> X -> E | 161.98 | 149.02 | 166.01 | 27348760.26 |
| B -> J -> E -> X | 139.12 | 163.55 | 170.17 | 27683632.54 |
| E -> X -> B -> J | 216.34 | 160.60 | 170.91 | 28227148.38 |
| E -> B -> X -> J | 216.34 | 131.63 | 157.55 | 28337516.04 |
| J -> E -> X -> B | 113.02 | 138.72 | 153.70 | 29712644.56 |
| J -> E -> B -> X | 193.47 | 146.16 | 161.70 | 30605810.78 |
| X -> J -> E -> B | 139.12 | 134.59 | 156.81 | 31884381.42 |
| E -> B -> J -> X | 242.44 | 156.46 | 174.02 | 32653825.73 |
| X -> E -> B -> J | 245.67 | 179.26 | 182.97 | 34259717.91 |
| E -> J -> B -> X | 190.24 | 152.33 | 166.12 | 34664297.1 |
| X -> J -> B -> E | 242.44 | 156.46 | 174.02 | 35353092.99 |
| J -> X -> E -> B | 165.22 | 142.85 | 161.60 | 35731859.84 |
| J -> B -> X -> E | 216.34 | 160.60 | 170.91 | 37273702.45 |
| E -> X -> J -> B | 161.98 | 149.02 | 166.01 | 38873791.13 |
| J -> B -> E -> X | 245.67 | 179.26 | 182.97 | 39535443.69 |
| X -> E -> J -> B | 139.12 | 163.55 | 170.17 | 39803700.42 |

**Supplementary Table 2. Cryo-EM data collection, refinement and validation statistics**

|  | **Smc5/6 Hexamer Apo State** |  | **Smc5/6 Hexamer Juxtaposed State** |  |  |  |  | **Smc5/6 Hexamer +ATP State** | **Smc5/6 Hexamer +ATP/DNA State** |
| --- | --- | --- | --- | --- | --- | --- | --- | --- | --- |
|  | **Apo Bridge** | **Apo Cross** | **Overall** | **Head** | **Opened Hinge** | **Intermediate Hinge** | **Closed Hinge** | **Head** | **Head** |
|  | **EMD-100183** | **EMD-100184** | **EMD-100185** | **EMD-100186** | **EMD-100187** | **EMD-100188** | **EMD-100189** | **EMD-100190** | **EMD-100191** |
|  | **34VB** | **34VC** | **34VD** | **34VE** |  |  |  | **34VF** | **34VG** |
|  |  |  | **Data collection and processing** |  |  |  |  |  |  |
| Magnification | 105,000 | 105,000 | 105,000 | 105,000 | 105,000 | 105,000 | 105,000 | 105,000 | 105,000 |
| Voltage (kV) | 300 | 300 | 300 | 300 | 300 | 300 | 300 | 300 | 300 |
| Total electron exposure (e<sup>-</sup>/Å<sup>2</sup>) | 60 | 60 | 60 | 60 | 60 | 60 | 60 | 60 | 60 |
| Defocus range (μm) | -1 to -2.5 | -1 to -2.5 | -1 to -2.5 | -1 to -2.5 | -1 to -2.5 | -1 to -2.5 | -1 to -2.5 | -1 to -2.5 | -1.3 to -2.5 |
| Voxel size (Å) | 0.831 | 0.831 | 0.837 | 0.837 | 0.837 | 0.837 | 0.837 | 0.831 | 0.837 |
| Symmetry imposed | C1 | C1 | C1 | C1 | C1 | C1 | C1 | C1 | C1 |
| Initial particle images (no.) | 572609 | 572609 | 810430 | 156402 | 153867 | 153867 | 153867 | 2408344 | 337576 |
| Final particle images (no.) | 78215 | 147532 | 26136 | 27702 | 43894 | 67772 | 26014 | 344402 | 84394 |
| Map resolution (Å) | 8.4 | 4 | 8.9 | 3.8 | 7.5 | 7.5 | 7.8 | 3.8 | 3.67 |
| FSC threshold | 0.143 | 0.143 | 0.143 | 0.143 | 0.143 | 0.143 | 0.143 | 0.143 | 0.143 |
|  | **Refinement** |  |  |  |  |  |  |  |  |
| Initial model used (PDB code) | 7QCD | 8WJN | 8I13 | 8WJN |  |  |  | 7TVE | 7TVE |
|  | R.m.s. deviations |  |  |  |  |  |  |  |  |
| Bond lengths (Å) | 0.005 | 0.004 | 0.004 | 0.004 |  |  |  | 0.004 | 0.004 |
| Bond angles (°) | 1.003 | 0.84 | 0.87 | 0.84 |  |  |  | 1.11 | 0.96 |
|  | Validation |  |  |  |  |  |  |  |  |
| MolProbity score | 1.43 | 1.54 | 1.1 | 1.54 |  |  |  | 2.63 | 2.38 |
| Clashscore | 0.87 | 0.78 | 0.62 | 0.78 |  |  |  | 20.08 | 11.5 |
| Poor rotamers (%) | 0 | 2.85 | 0 | 2.85 |  |  |  | 2.94 | 4.37 |
|  | Ramachandran plot |  |  |  |  |  |  |  |  |
| Favored (%) | 84.83 | 92.9 | 94.11 | 92.9 |  |  |  | 92.22 | 95.39 |
| Allowed (%) | 14.87 | 7.1 | 5.89 | 7.1 |  |  |  | 7.6 | 4.37 |
| Disallowed (%) | 0.3 | 0 | 0 | 0 |  |  |  | 0.17 | 0.24 |

### **Supplementary Movie Captions**

**Supplementary Movie1: CryoDRGN of Cross state.** Volumes reconstructed along the first principal component (PC1) of the CryoDRGN latent space learned from the Cross state particles show the Smc6 head movement.

**Supplementary Movie2: 3DVA of the Juxtaposed state head.** Volumes reconstructed along the first principal component (PC1) identified by 3D variability analysis (3DVA) in cryoSPARC for particles assigned to Juxtaposed state show the Smc6 head movement.

**Supplementary Movie3: Model of the proposed pre-catalytic activation pathway.** ChimeraX-generated morph illustrating transitions between the four states arranged according to the proposed pathway.

### **Supplementary Figures**

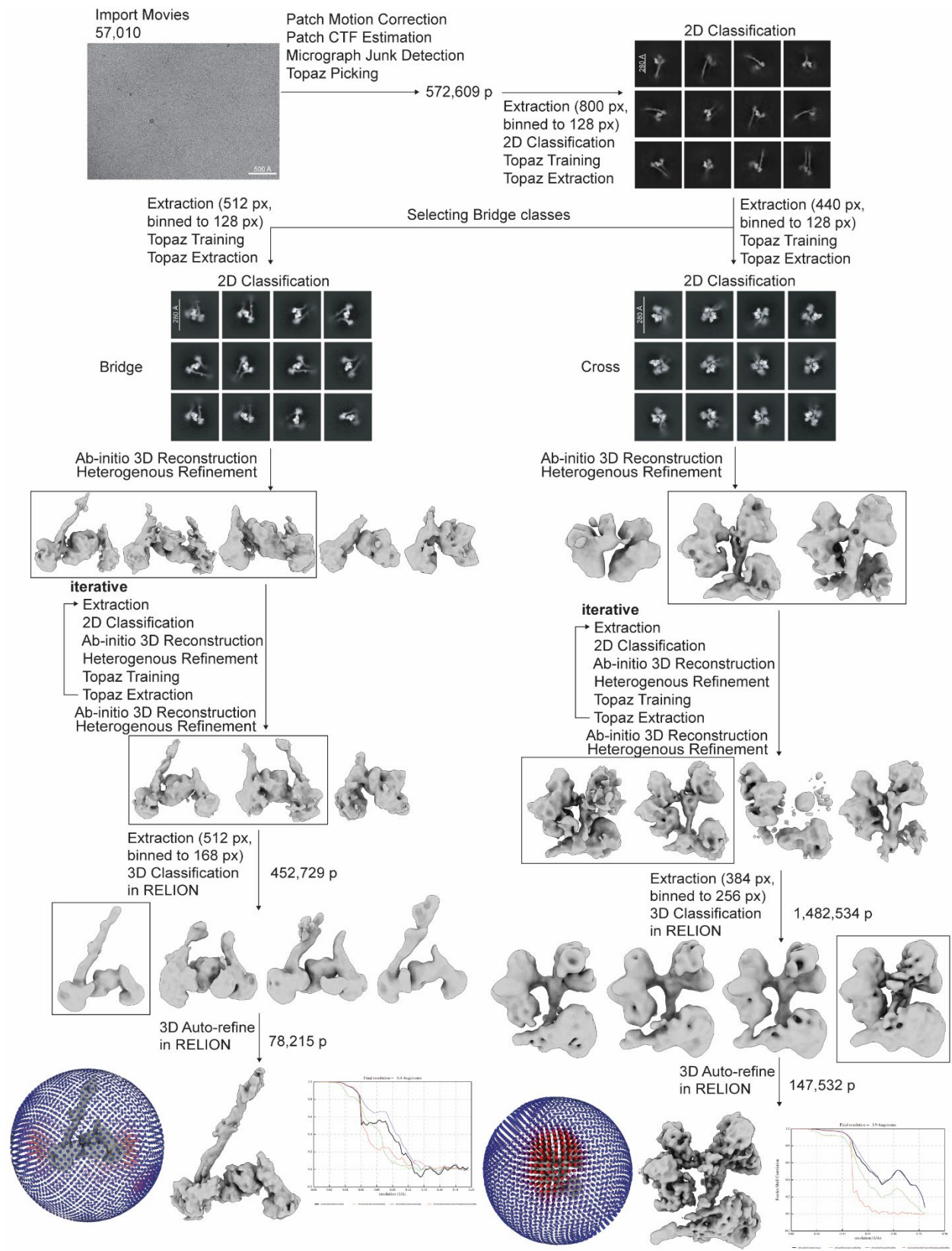

**Supplementary Figure 1: CryoEM data processing workflow of Bridge and Cross state**

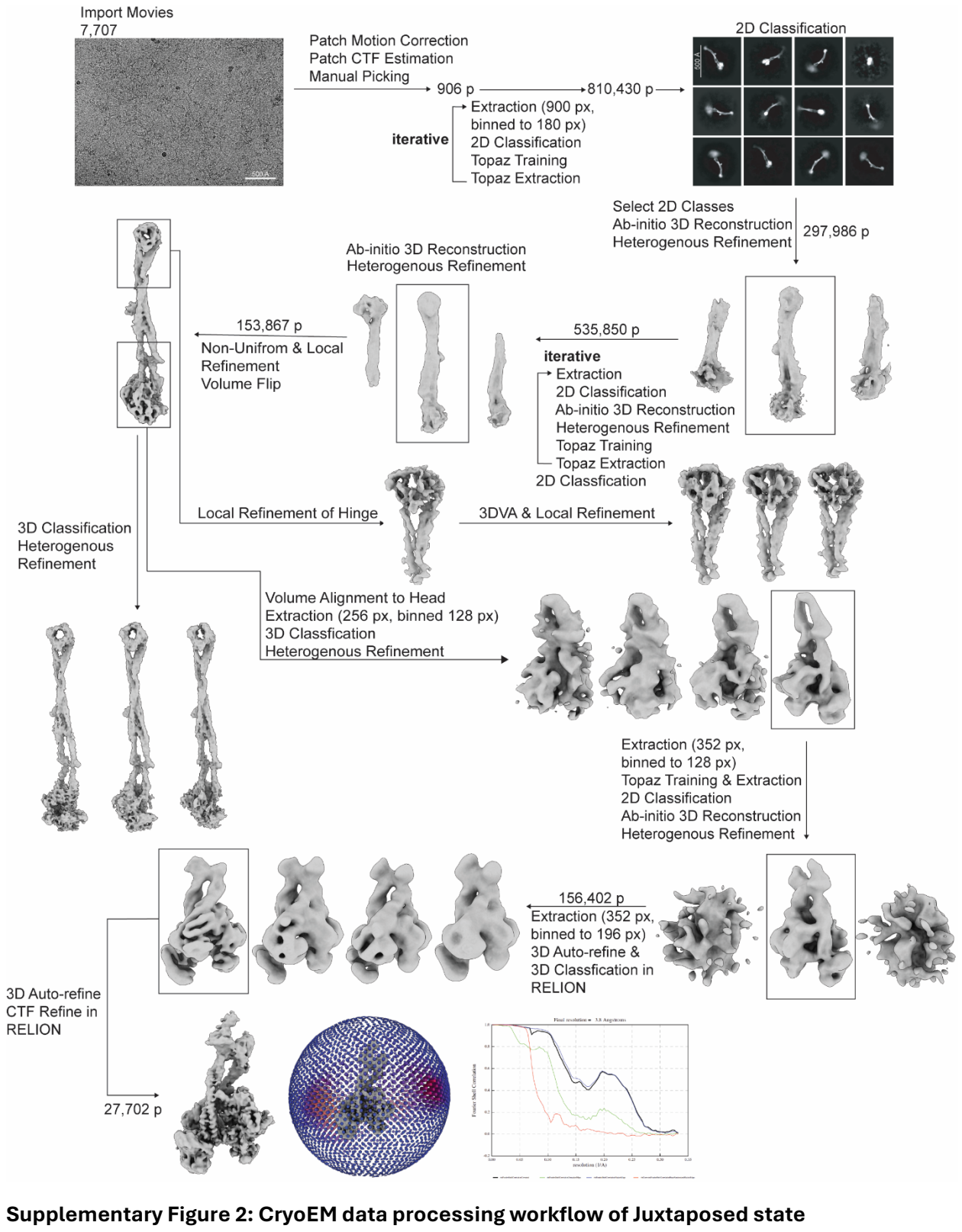

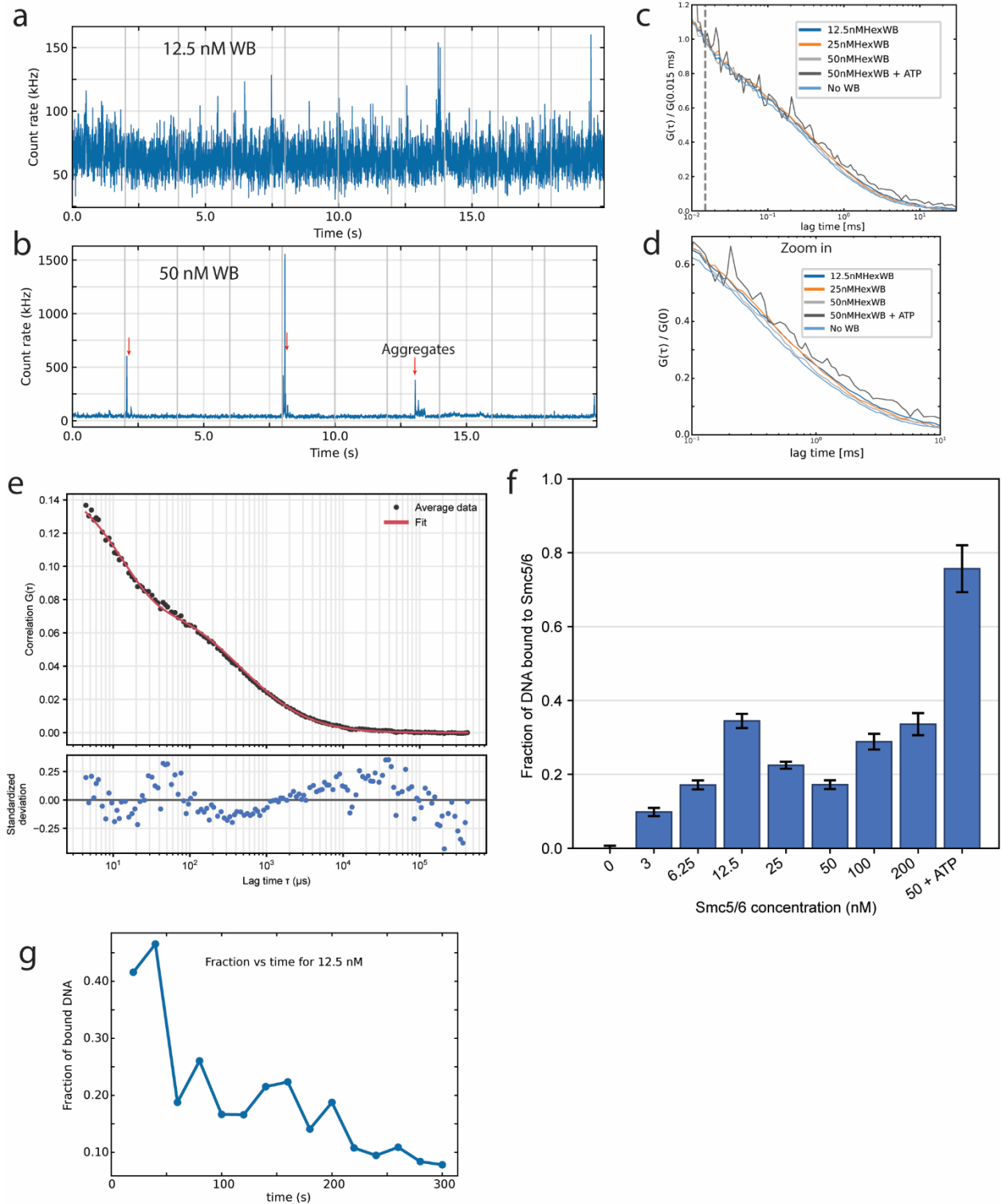

**Supplementary Figure 3: FCS analysis of DNA interactions**

FCS of 25 nM Cy5 labeled 72 bp dsDNA in the presence of increased concentrations of the Smc5/6 Waker B mutant. (a, b) Representative intensity time traces at (a) 12.5 nM and (b) 50 nM Smc5/6. At higher concentration, large intensity bursts appear (red arrows) corresponding to protein-DNA

aggregates. The section containing these bursts were discarded from further FCS fitting. (c) Autocorrelation curves $G(\tau)$ at different concentrations, normalized to $G(\tau)$ at 15 μs (dashed line). The shift of the decay towards the right with increasing Smc5/6 concentration reflects slower DNA diffusion upon binding. Addition of ATP at 50 nM Smc5/6 leads to the largest shift while the control without Smc5/6 makes free DNA base line. (d) Zoom in of the correlation curves highlighting the shift in the diffusion times. (e) Example of fitting and residual plot of the autocorrelation curve at 12.5 nM with a two diffusion and one triplet component model. The bound fractions in f and g are extracted from this fit. (f) Fraction of DNA bound to Smc5/6 extracted from the fitting of two component diffusion as a function of Smc5/6 concentration. The bound fraction increases upto ∼12.5 nM and then goes down likely due to the aggregation as noticed in the intensity traces. (g) Bound fraction at 12.5 nM Smc5/6 as a function of time from the start of the measurement showing a decrease of over the 300s measurement showing the rapid dissociation of Smc5/6 at room temperature.

**a**

Cross 3DVA

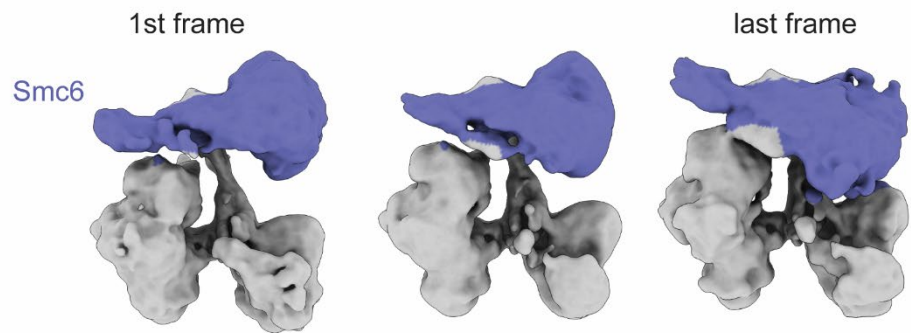

**b**

Cross CryoDRGN

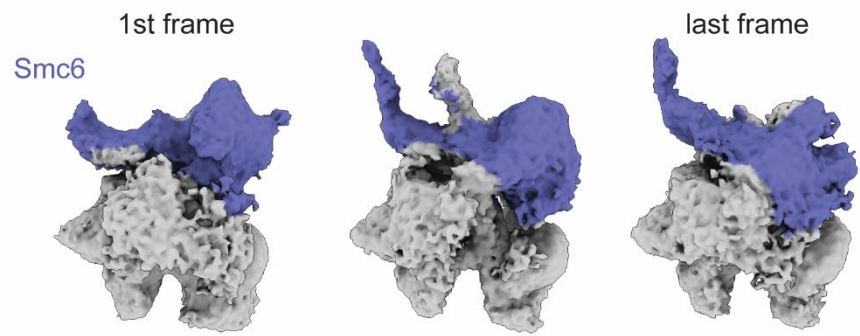

**c**

Juxtaposed 3DVA

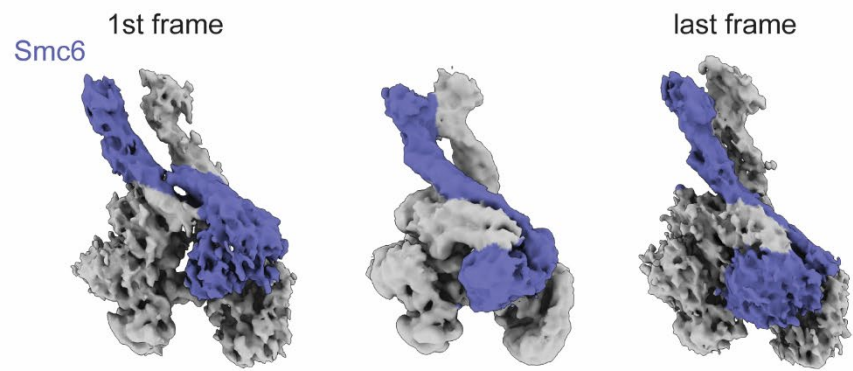

**d**

Juxtaposed CryoDRGN

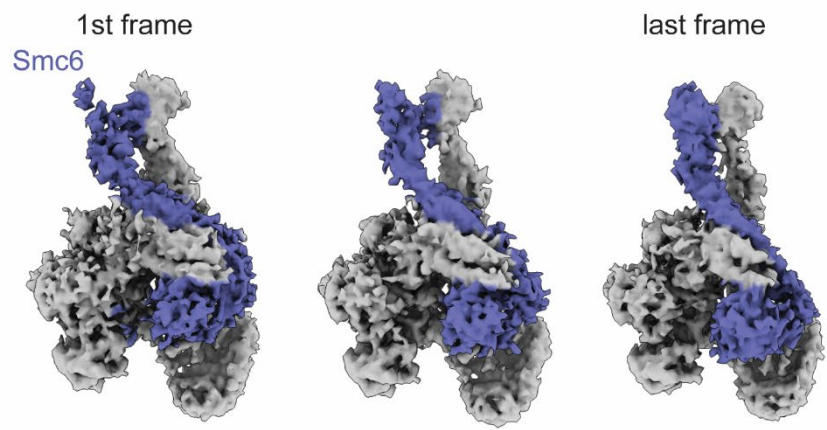

**Supplementary Figure 4: 3DVA and CryoDRGN analysis of Cross and Juxtaposed state.**

Heterogenous analysis of Cross (a,b) and Juxtaposed (c, d) state using 3DVA (a, c) and CryoDRGN (b, d) showing the first (left), intermediate (middle) and last frame of the principle components that show Smc6 movement. Both 3DVA and CryoDRGN analysis shows the Smc6 (Blue) head migrates to Smc5 in both Cross and Juxtaposed state. CryoDRGN analysis resolved the bending of Smc6 joint in the Cross state.

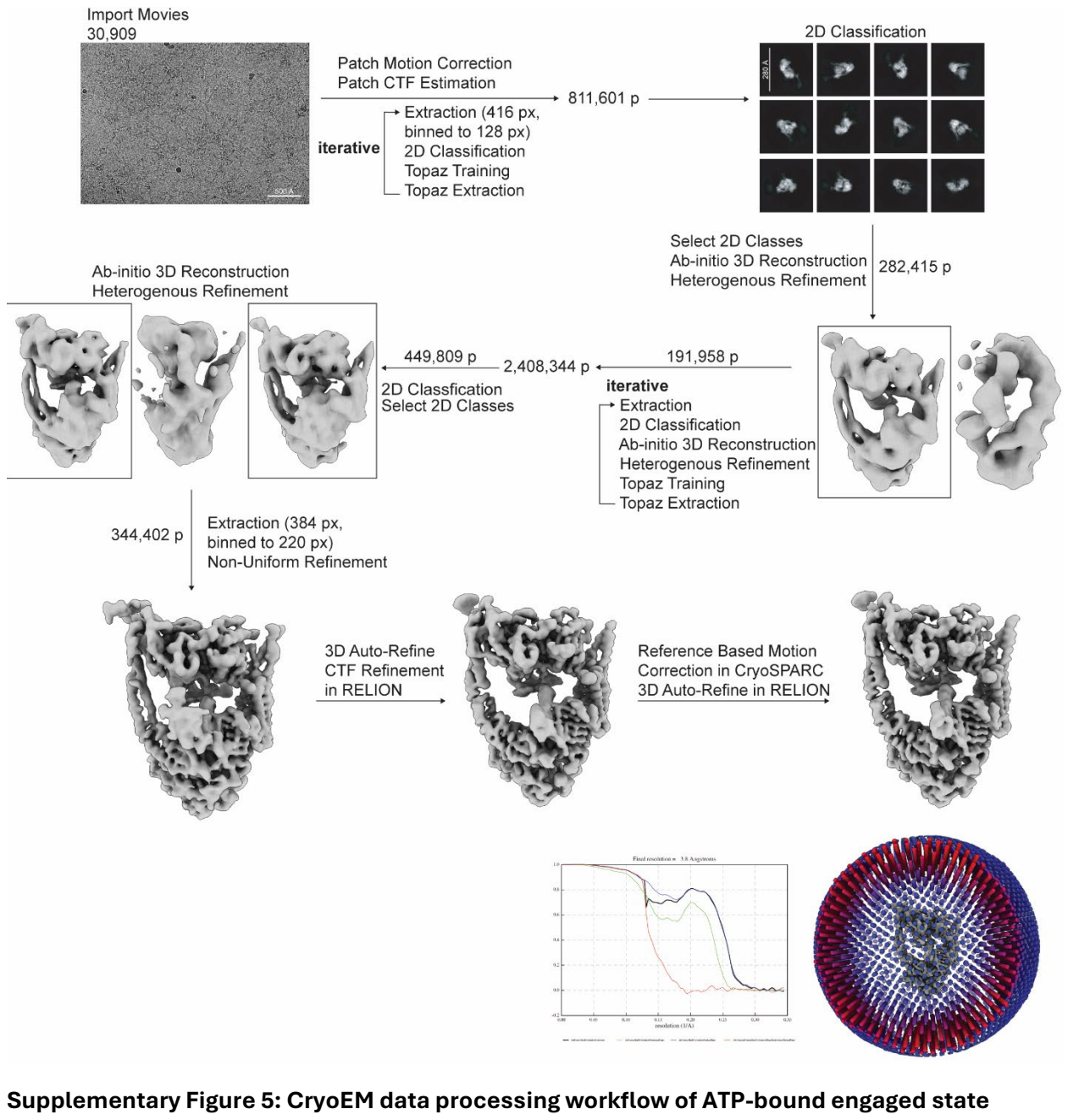

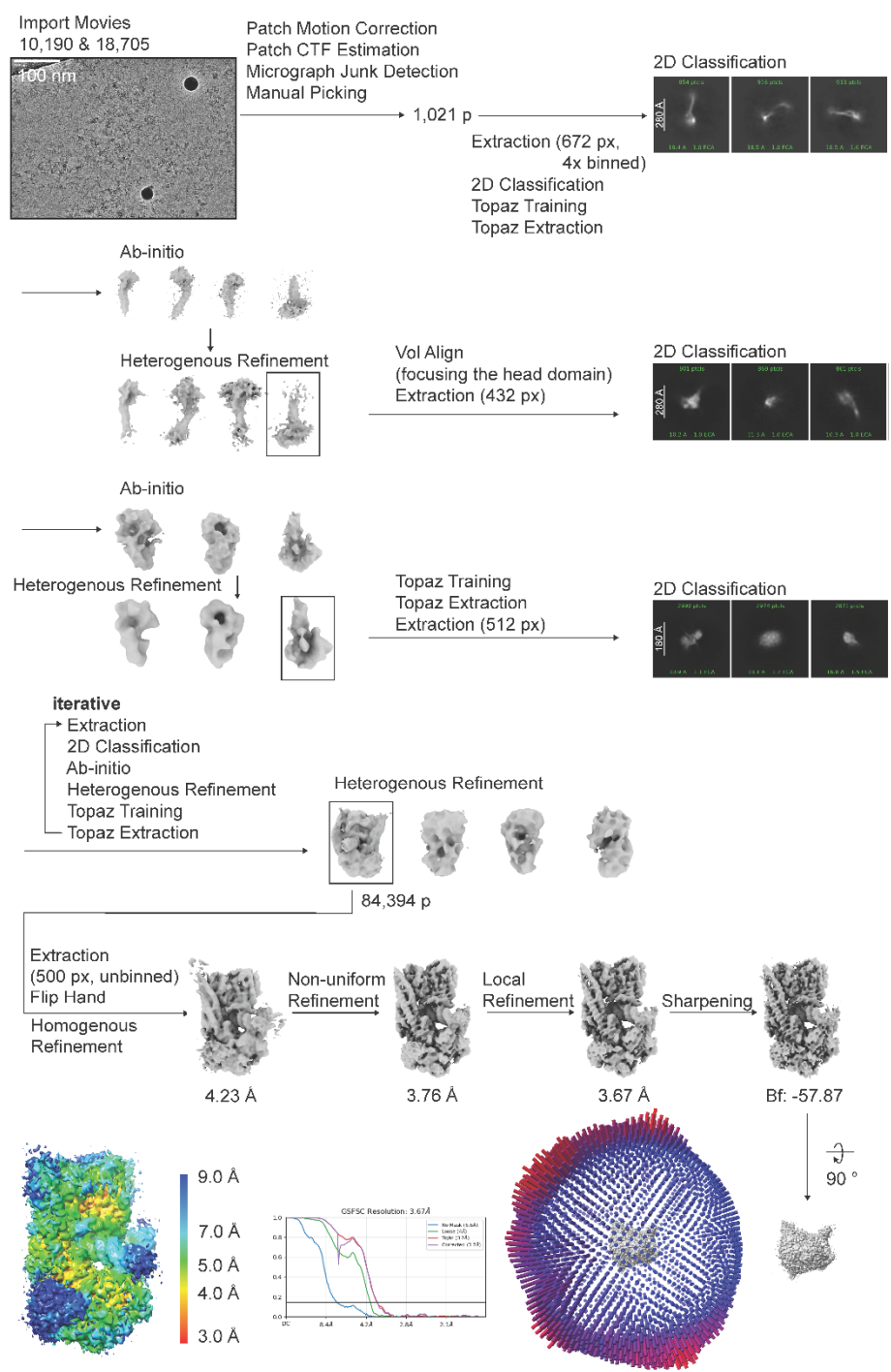

**Supplementary Figure 6: CryoEM data processing workflow of ATP/DNA-bound engaged state**

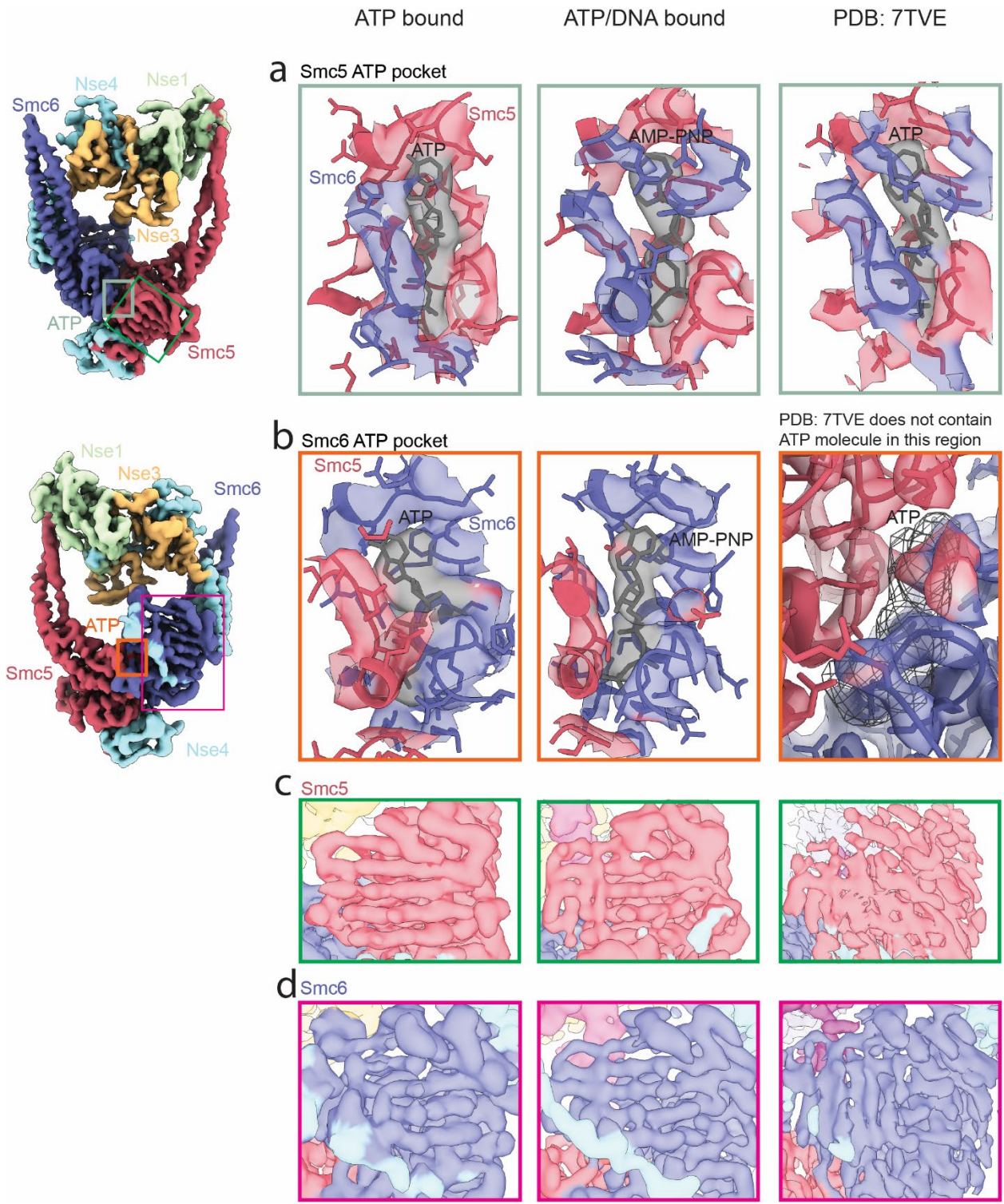

**Supplementary Figure 7: ATP pockets and better resolved beta strands from Smc5 and Smc6.**
a, ATP pocket in Smc5. b, ATP pocket in Smc6. PDB: 7TVE (right) does not contain ATP molecule in this region. Mesh is used to show the possible ATP location. c, Beta sheet in Smc5 ATPase domain. d, Beta sheet in Smc6 ATPase domain.

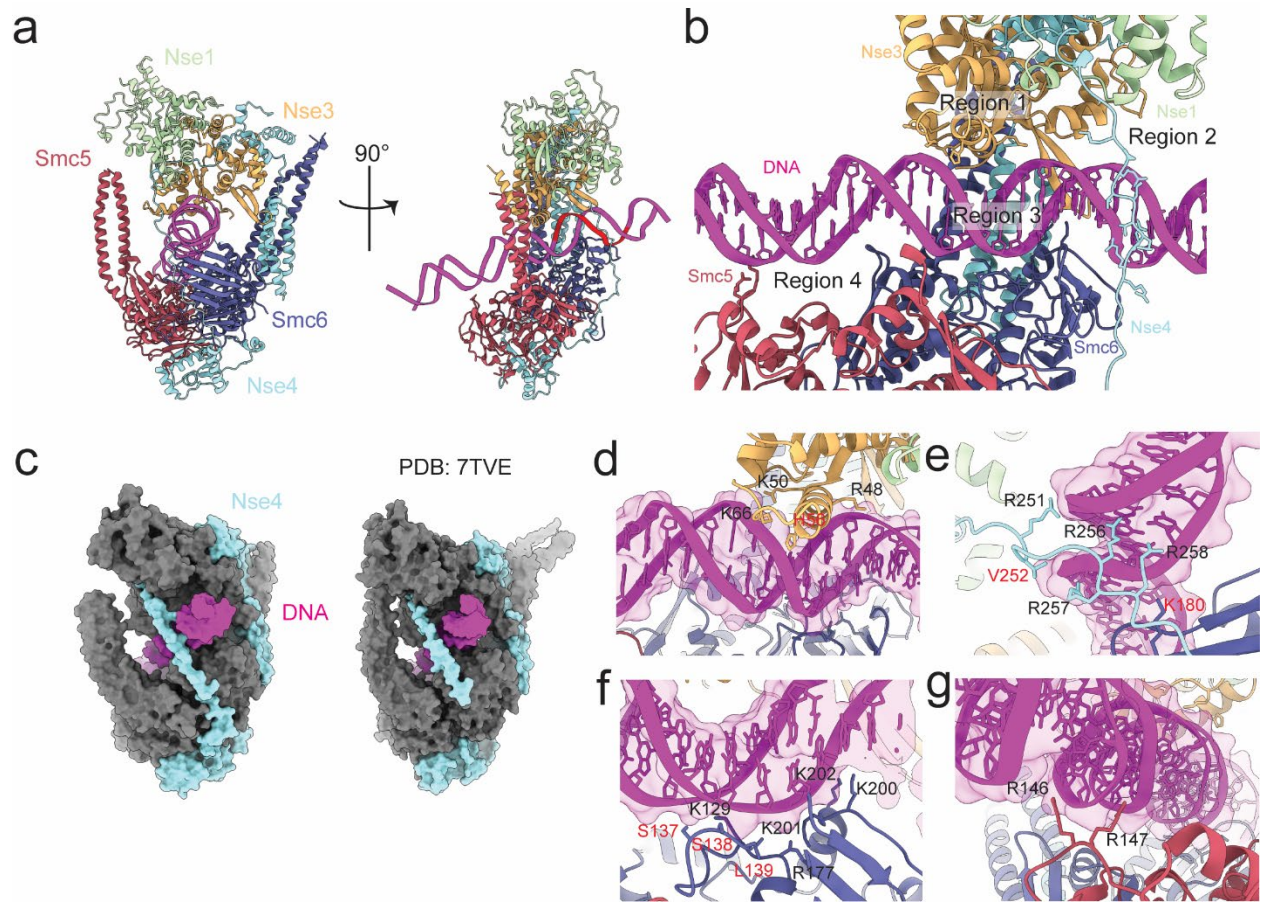

**Supplementary Figure 8: Smc5/6 interaction with DNA strand** a, The DNA strand identified within the binding pocket, shown in red. b, Close-up view of the DNA within the binding cavity of Smc5/6, with the interacting residues are grouped into four distinct regions. c, Comparison of Nse4 loop encasing the DNA in the present model and in 7TVE. Region 1 (d), 2 (e), Region 3 (f), and Region 4 (g) of Smc5/6 residues interacting with the DNA. Newly identified binding residues are marked as red.
